# Phenotypic Screening Identifies Small-Molecule Inhibitors with Distinct Activities across the BK Polyomavirus Life Cycle

**DOI:** 10.64898/2026.08.20.745923

**Authors:** Clara Husser, Hannes Roggenkamp, Emma Kraus, Patrick Blümke, Sanamjeet Virdi, Claudia Schmidt, Jessica Rückert, Thomas Schulz, Adam Grundhoff, Nicole Fischer

## Abstract

**BACKGROUND:** BK polyomavirus (BKPyV) reactivation is a major complication in kidney and hematopoietic stem cell transplant recipients, yet no specific antiviral therapy is currently available. Antiviral discovery is complicated by the restricted tropism and slow replication kinetics of BKPyV and its extensive dependence on cellular processes.

**RESULTS:** We established a phenotypic high-throughput screening and validation pipeline to identify small-molecule inhibitors of BKPyV infection. Using an SV40-infected CV-1 reporter system, approximately 28,000 small molecules were screened, yielding 98 primary candidates. Confirmatory testing identified 33 compounds with reproducible activity, of which 16 subsequently inhibited BKPyV in human renal proximal tubular epithelial cells. Concentration–response and cytotoxicity analyses revealed distinct antiviral potency and selectivity profiles, and integration of these data with predicted toxicity, physicochemical properties, and synthetic accessibility enabled further compound prioritization. Time-of-addition experiments revealed distinct temporal windows of antiviral activity, and MOI-dependent concentration–response analyses demonstrated that the potency of selected inhibitors varied with viral inoculum. Further characterization of prioritized compounds identified differential effects on BKPyV attachment and viral gene expression. Transcriptomic profiling of three selected compounds C5, C8, and C9 revealed distinct compound-associated cellular responses, supporting interference with different host-dependent processes during BKPyV infection.

**CONCLUSIONS:** We identified a pharmacologically diverse panel of small-molecule inhibitors active against BKPyV in human renal epithelial cells. Their distinct potency, selectivity, temporal activity, and cellular response profiles indicate multiple modes of antiviral interference and establish C5, C8, and C9 as candidates for further target identification and optimization. More broadly, our findings demonstrate the utility of surrogate phenotypic screening for discovering inhibitors of BKPyV and provide new chemical tools to investigate host dependencies of the BKPyV life cycle.

## Background

Human polyomaviruses are widespread, small, non-enveloped DNA viruses that establish lifelong persistent infections in the majority of the human population (1). Although primary infection is typically asymptomatic, four human polyomaviruses are associated with severe disease, predominantly under conditions of immunosuppression. Merkel cell polyomavirus (MCPyV) is causally linked to Merkel cell carcinoma (2), an aggressive skin cancer associated with high mortality. Trichodysplasia spinulosa polyomavirus (TSPyV) causes uncontrolled viral replication in hair follicles of immunosuppressed patients, resulting in the characteristic skin disease trichodysplasia spinulosa (3). JC polyomavirus (JCPyV) causes progressive multifocal leukoencephalopathy (PML), a rapidly progressive and frequently fatal demyelinating disease of the central nervous system (1, 4). BK polyomavirus (BKPyV), in turn, is a major cause of polyomavirus-associated nephropathy (PVAN) in kidney transplant recipients and polyomavirus-associated hemorrhagic cystitis (PVHC) in hematopoietic stem cell transplant recipients (1, 5).

BKPyV- and JCPyV-associated diseases represent a substantial clinical burden, particularly among transplant recipients and other immunocompromised populations. JCPyV-associated PML is associated with mortality rates exceeding 50%, with a particularly high incidence observed in patients with multiple sclerosis receiving integrin antagonist therapies (6–8). BKPyV reactivation occurs frequently in immunosuppressed kidney transplant recipients: approximately 10–30% develop BKPyV DNAemia, and 1–10% progress to BKPyV-associated nephropathy, which can result in graft dysfunction and ultimately graft loss (9–13). Given the large number of kidney and hematopoietic stem cell transplantations performed worldwide, BKPyV-associated disease remains an important complication of transplantation. Despite this considerable clinical impact, no virus-specific antiviral therapy is currently approved for the treatment or prevention of BKPyV or JCPyV infection.

A VP1-specific human monoclonal neutralizing antibody with activity against all major BKPyV serotypes has recently been reported (14) and is currently being evaluated in a randomized phase II/III clinical trial (15). Nevertheless, current management of BKPyV infection continues to rely predominantly on reduction of immunosuppressive therapy to restore virus-specific immune control (16–18). Although reduction of immunosuppression can facilitate viral clearance in kidney transplant recipients, it increases the risk of acute rejection and may ultimately compromise graft function or survival. Therapeutic options are even more limited for BKPyV-associated hemorrhagic cystitis after hematopoietic stem-cell transplantation, where management remains largely supportive. Several compounds, including cidofovir, leflunomide, and mefloquine, have been evaluated as potential treatment options; however, controlled clinical studies have failed to demonstrate consistent antiviral efficacy, and toxicity, including nephrotoxicity, can further limit their clinical utility (19, 20). Other approaches, including immunomodulatory drugs and antibiotics, have likewise shown limited or inconsistent benefit in clinical trials (21). Thus, there remains a substantial unmet need for antiviral agents that directly and selectively interfere with BKPyV infection.

The identification of such antiviral agents is complicated by the biology of polyomaviruses. Their small genomes encode only a limited repertoire of viral proteins, requiring extensive exploitation of cellular proteins and pathways to complete the viral replication cycle. This dependence on host-cell functions provides opportunities to pharmacologically interfere with cellular processes required for productive infection and may offer an alternative to conventional strategies directed exclusively against virally encoded proteins. At the same time, however, antiviral discovery for BKPyV has been hindered by their restricted host and cell tropism, slow replication kinetics, and lack of a prominent cytopathic effect *in vitro*. Together, these characteristics complicate the establishment of robust, scalable screening systems suitable for systematic antiviral compound discovery.

Specifically, BKPyV is a small, non-enveloped virus with a circular double-stranded DNA genome of approximately 5 kb. The viral genome is organized into early and late transcriptional regions separated by a non-coding control region (NCCR), which contains the viral origin of replication and bidirectional promoters controlling viral gene expression (1). Infection is initiated by attachment to sialylated ganglioside receptors at the cell surface, followed by endocytic uptake and retrograde trafficking through intracellular compartments to the endoplasmic reticulum (22–25). Productive infection requires extensive engagement of cellular pathways during subsequent steps, including chaperone-assisted capsid disassembly, delivery of the viral genome to the nucleus, and recruitment of the cellular machinery required for viral DNA replication. Early viral gene expression produces, most prominently, the large T antigen (LT) and small t antigen (sT), which modulate cellular pathways and promote a cellular environment permissive for viral genome replication (1). In addition to its protein-coding genes, BKPyV expresses a virus-derived microRNA during infection that contributes to post-transcriptional regulation of viral gene expression and modulation of host immune recognition (26, 27).

Following viral genome replication, activation of the late transcriptional program results in expression of the structural capsid proteins VP1, VP2, and VP3, as well as the auxiliary agnoprotein (1, 28, 29). Viral capsid assembly occurs in the nucleus and ultimately culminates in the production and release of progeny virions. Thus, productive BKPyV infection requires the coordinated progression through attachment, entry and intracellular trafficking, early viral gene expression, genome replication, late gene expression, virion assembly, and release. Throughout these stages, BKPyV remains highly dependent on cellular functions, notably because the virus does not encode its own DNA polymerase and therefore relies on the host-cell replication machinery for amplification of its genome.

Despite this general understanding of the BKPyV life cycle, important aspects of viral replication and the cellular processes supporting individual stages of infection remain incompletely understood. Mechanistic insights into polyomavirus entry and replication have frequently been obtained using surrogate polyomavirus models or transformed cell lines (22, 24), whereas studies performed directly in primary human target cells remain comparatively limited. This restricted experimental accessibility has complicated systematic analysis of the cellular processes required at different temporal stages of BKPyV infection. It has similarly constrained the development of scalable antiviral screening approaches capable of identifying pharmacological vulnerabilities throughout the viral life cycle.

Phenotypic screening provides one strategy to overcome some of these limitations. In contrast to target-based approaches, phenotypic screening identifies compounds on the basis of their ability to alter an infection-associated phenotype without requiring prior knowledge of the molecular target. This approach is particularly attractive for viruses such as BKPyV, which encode relatively few proteins and depend extensively on cellular functions throughout their replication cycle. However, the biological characteristics of BKPyV itself, including its restricted tropism, slow replication kinetics, and limited cytopathic effect, make direct high-throughput screening technically challenging.

To address these limitations, we employed a phenotypic high-throughput screening strategy using simian virus 40 (SV40), a closely related polyomavirus, as a surrogate screening model in CV-1 reporter cells under biosafety level 2 conditions. The surrogate system enabled scalable identification of compounds capable of interfering with productive polyomavirus infection without prespecifying a viral or cellular molecular target. Compounds identified in the SV40 screen were subsequently evaluated for antiviral activity against BKPyV in human renal proximal tubular epithelial cells, representing the relevant target cell type for BKPyV infection. This sequential strategy combined the scalability of a surrogate phenotypic screening system with validation against the clinically relevant virus in human renal epithelial cells.

In this study, we used this screening and validation pipeline to identify and pharmacologically characterize small-molecule inhibitors of BKPyV infection. Following confirmation of antiviral activity, compounds were evaluated for concentration-dependent inhibition, cytotoxicity, selectivity, and temporal profiles of antiviral activity, enabling prioritization of compounds with distinct antiviral phenotypes. Selected compounds were subsequently subjected to further functional and transcriptomic characterization to investigate their effects on different stages of BKPyV infection and the cellular responses associated with compound treatment. Together, this approach identifies chemically and functionally distinct inhibitors of BKPyV infection and provides pharmacological tools for investigating host-dependent processes required for productive polyomavirus infection.

## Results

### Phenotypic screening identifies small-molecule inhibitors with activity against BKPyV

To identify small molecules with antiviral activity against polyomaviruses, we established a phenotypic screening cascade using SV40 as a surrogate virus, followed by validation against BKPyV in human renal epithelial cells. CV-1 reporter cells constitutively expressing red fluorescent protein (RFP) under a cellular promoter were used for the primary screen. CV-1 reporter cells were infected with SV40 at an MOI of 0.5 and treated with the compounds at 10 µM concentration. Productive SV40 infection resulted in a cytopathic phenotype accompanied by loss of the constitutively expressed RFP signal, whereas inhibition of infection preserved reporter fluorescence (Fig. 1A). This phenotypic approach enabled target-agnostic identification of antiviral compounds while inherently counterselecting against compounds with pronounced cytotoxicity in CV-1 cells. Prior to compound screening, the assay system was evaluated using two previously described inhibitors of polyomavirus replication with distinct antiviral properties. Cidofovir, an acyclic nucleoside phosphonate with established inhibitory activity against polyomavirus DNA replication, was included as a reference antiviral compound, whereas hexachlorophene has previously been identified as an inhibitor of the ATPase activity of polyomavirus large T antigen and shown to suppress both SV40 and BKPyV replication (30, 31). Based on its robust activity in the reporter assay (suppl. Fig.1) and reduced cytotoxicity compared to Cidofovir, hexachlorophene was selected as the positive control for the screening and included in multiple wells on each screening plate to monitor assay performance and facilitate plate-to-plate calibration. Antiviral activity was assessed using two independent readouts, plate reader-based quantification of RFP fluorescence and fluorescence microscopy, which showed a high degree of concordance (suppl. Fig. S1B-C, Fig. 1B–C). Primary hit selection was performed on a plate-by-plate basis using two independent selection criteria: a relative RFP fluorescence intensity (FI) >20% or a >2-fold increase in relative FI. Representative screening plates and the corresponding plate reader- and imaging-based hit selection are shown in Suppl. Fig. S2. Application of these criteria identified 98 primary hits, which were subsequently retested in a confirmatory screen using independent replicate measurements under the same experimental conditions. Antiviral activity was assessed independently by plate reader-based quantification of RFP fluorescence and fluorescence microscopy, with both readouts showing a high degree of concordance (Fig. 1B–C). Compounds were considered confirmed hits and selected for downstream analysis if their relative RFP fluorescence intensity reached at least 50% of that observed with the positive-control compound hexachlorophene. Based on this predefined selection criterion, 33 of the 98 compounds were confirmed as active against the SV40-induced phenotype and advanced to subsequent validation against BKPyV (Fig. 1C).

**Figure 1.**
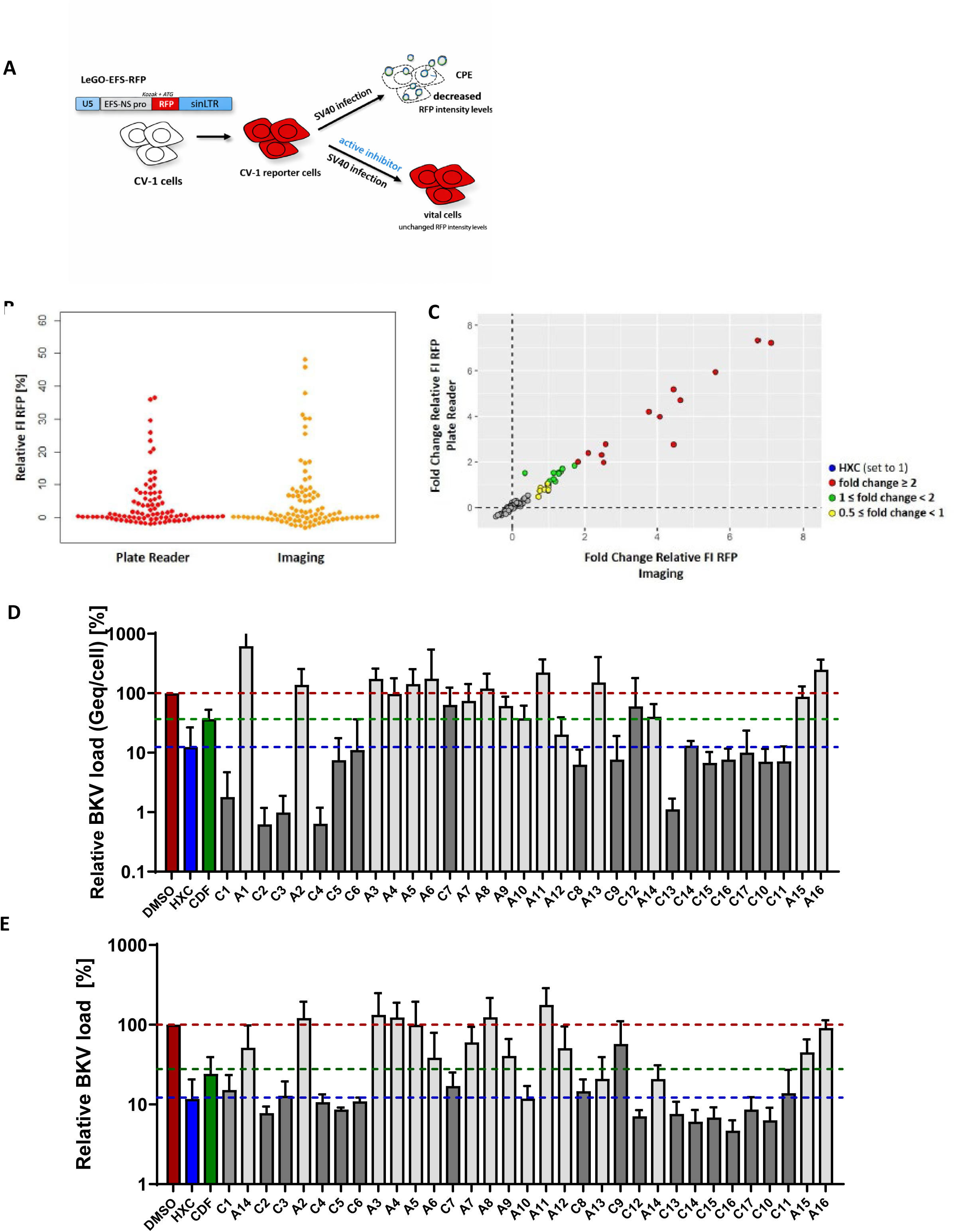
Phenotypic screening strategy and summary of results from the surrogate and BKPyV-specific screens. **(A)** Schematic of the phenotypic cell-based screen used to identify inhibitors of SV40 and the related polyomaviruses BKPyV and JCPyV. African green monkey CV-1 cells stably expressing red fluorescent protein (RFP) from the elongation factor 1α (EF1α) promoter were generated by lentiviral transduction followed by neomycin selection. Infection with SV40 results in efficient viral replication and cytopathic effect (CPE), leading to a reduction in RFP signal. Antiviral compounds prevent or reduce virus-induced CPE, which is quantified by measuring RFP fluorescence intensity. **(B, C)** Thirty-three of 98 compounds were confirmed to exhibit strong inhibitory activity against SV40 in CV-1 reporter cells. The fluorescence intensity (FI) of uninfected cells was defined as 100%, whereas the FI of infected, DMSO-treated cells was defined as 0%. FI values for infected cells treated with HXC or candidate compounds were normalized to these controls. The FI of HXC-treated infected cells was set to 1, and the relative FI of candidate compounds was expressed as fold change relative to HXC. RFP fluorescence was quantified by high-content imaging or plate reader analysis, as indicated. The correlation between relative FI values obtained in the primary high-throughput screen (HTS) and those from the confirmatory screen for the 98 hit compounds is shown. **(D, E)** Sixteen of the 33 SV40-active compounds exhibited inhibitory activity against BKPyV infection. Human primary renal tubular epithelial cells (hRPTEC) were infected with BKPyV in the presence of DMSO, HXC, CDF, or one of the 33 candidate compounds. At 6 days post infection (p.i.), genomic DNA (gDNA) was isolated from cell pellets and culture supernatants, and viral genome equivalents (Geq) were quantified by qPCR (n = 6). Intracellular BKPyV load was determined by normalizing cellular Geq values to GAPDH (D), whereas extracellular BKPyV load was calculated from Geq values obtained from culture supernatants (E). BKPyV loads in infected, DMSO-treated cells were set to 100%.

Applying this selection criterion resulted in 33 confirmed hits, which were subsequently advanced to secondary screening against the clinically relevant BKPyV. To determine whether antiviral activity identified in the SV40 surrogate system translated to BKPyV, these compounds were evaluated in primary human renal proximal tubular epithelial (hRPTE) cells. Cells were infected with BKPyV at an MOI of 0.5 and treated with each compound at a concentration of 10 µM. At 6 days post-infection, viral genome equivalents were quantified by qPCR targeting the VP1 region in both cell-associated and culture supernatant fractions, thereby assessing effects on intracellular viral loads and progeny virus release, respectively. Compounds were considered active against BKPyV if both cell-associated and extracellular viral genome equivalents were reduced to ≤50% of the levels detected in DMSO-treated infected controls. Based on this predefined criterion, 16 of the 33 SV40-active compounds were confirmed as active against BKPyV (Fig. 1D–E), corresponding to a confirmation rate of approximately 48%. Sixteen of the 33 SV40-active compounds reduced BKPyV loads compared with DMSO-treated infected controls (Fig. 1D–E), corresponding to a confirmation rate of approximately 48% among the SV40-active compounds.

These results established a panel of 16 compounds with antiviral activity against BKPyV in its relevant human renal target cell type for subsequent pharmacological characterization.

### Dose–response and cytotoxicity profiling identifies BKPyV inhibitors with distinct selectivity profiles

To characterize antiviral potency and distinguish antiviral activity from nonspecific cytotoxicity, the 16 BKPyV-active compounds were evaluated in parallel concentration–response and cell viability assays. Primary hRPTE cells were infected with BKPyV at an MOI of 0.5 and treated with increasing concentrations of compounds C1–C16 (0.1 – 30µM). Extracellular BKPyV DNA was quantified in culture supernatants at 6 days post-infection, and concentration–response curves were used to derive approximate IC₅₀ values (Fig. 2). In parallel, cell viability was assessed in uninfected cells exposed to the compounds at concentrations ranging from 0.1 to 100 µM; after 6 days, cytotoxicity profiles were evaluated and CC₅₀ values were determined.

**Figure 2.**
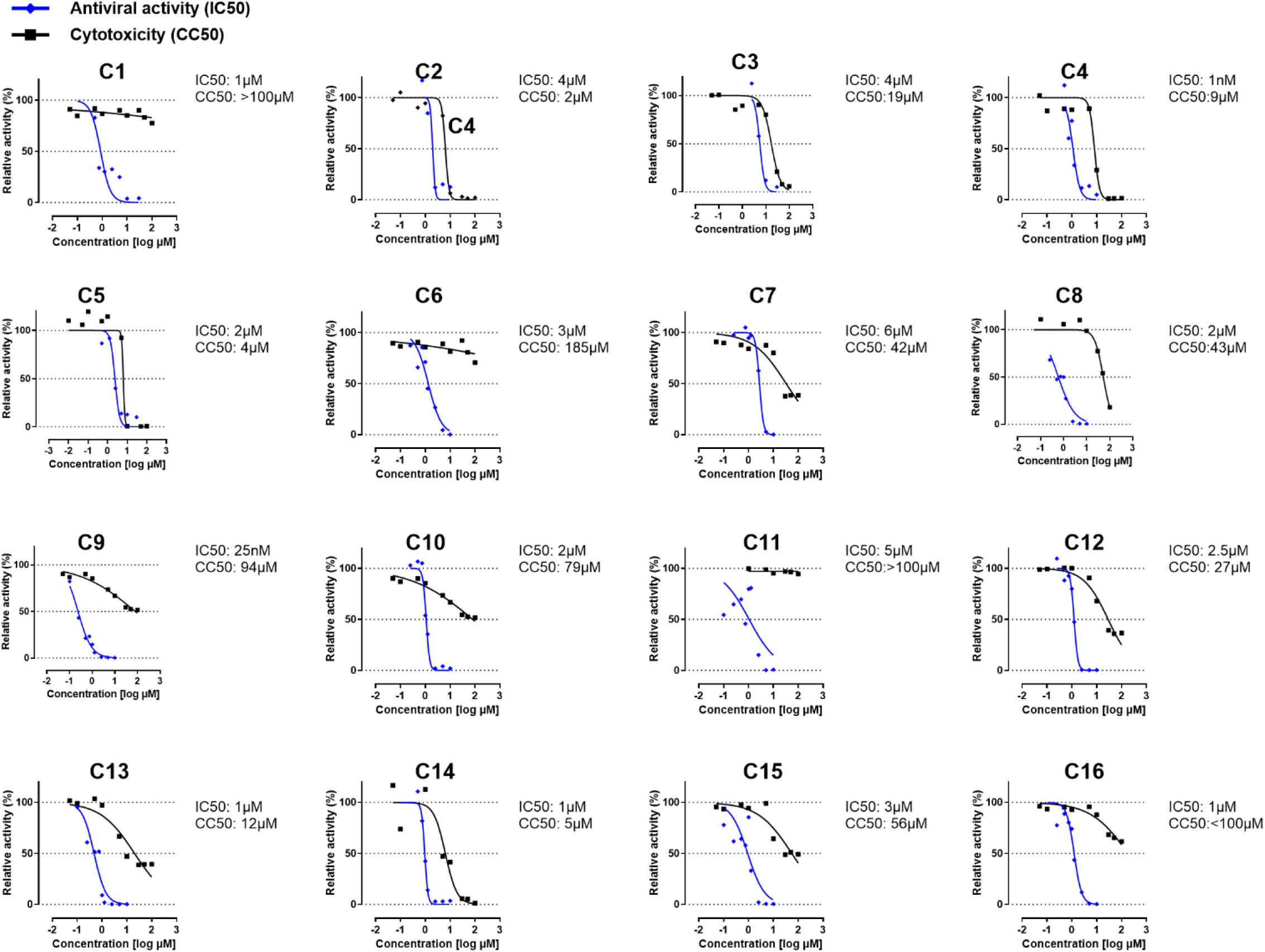
Antiviral activity and cytotoxicity of compounds C1–C16 in BKPyV-infected hRPTE cells. Primary human renal proximal tubular epithelial (hRPTE) cells were infected with BKPyV at a multiplicity of infection (MOI) of 0.5 in the presence of increasing concentrations of compounds C1– C16. At 6 days post infection (p.i.), viral replication was quantified by measuring BKPyV genomic DNA (gDNA) in culture supernatants using qPCR (n = 3). Viral loads in infected, DMSO-treated control cells were normalized to 100%. Dose–response curves (blue lines) and corresponding approximate IC₅₀ values are shown for each compound. In parallel, primary hRPTE cells were treated with increasing concentrations of compounds C1–C16 in the absence of infection to assess cytotoxicity. At 6 days post-treatment, cell viability was determined by MTT assay (n = 3), and results are shown as black lines.

The compounds displayed heterogeneous concentration–response profiles, with marked differences in both antiviral potency and cytotoxicity. To assess the separation between antiviral activity and cytotoxicity, selectivity indices (SI) were calculated as the ratio of CC₅₀ to IC₅₀. Eleven compounds (C1, C5, C6, C8-13, C15, and C16) exhibited SI values >10, indicating a greater than tenfold separation between antiviral activity and cytotoxicity under the conditions tested (Table 1). For subsequent experiments, C5, C8, and C9 were reordered after depletion of the initially available compound material. Reassessment of their concentration–response and cytotoxicity profiles with the newly obtained batches resulted in IC₅₀, CC₅₀, and consequently SI values that differed from those obtained with the original material (Table 1).

**Table 1:**
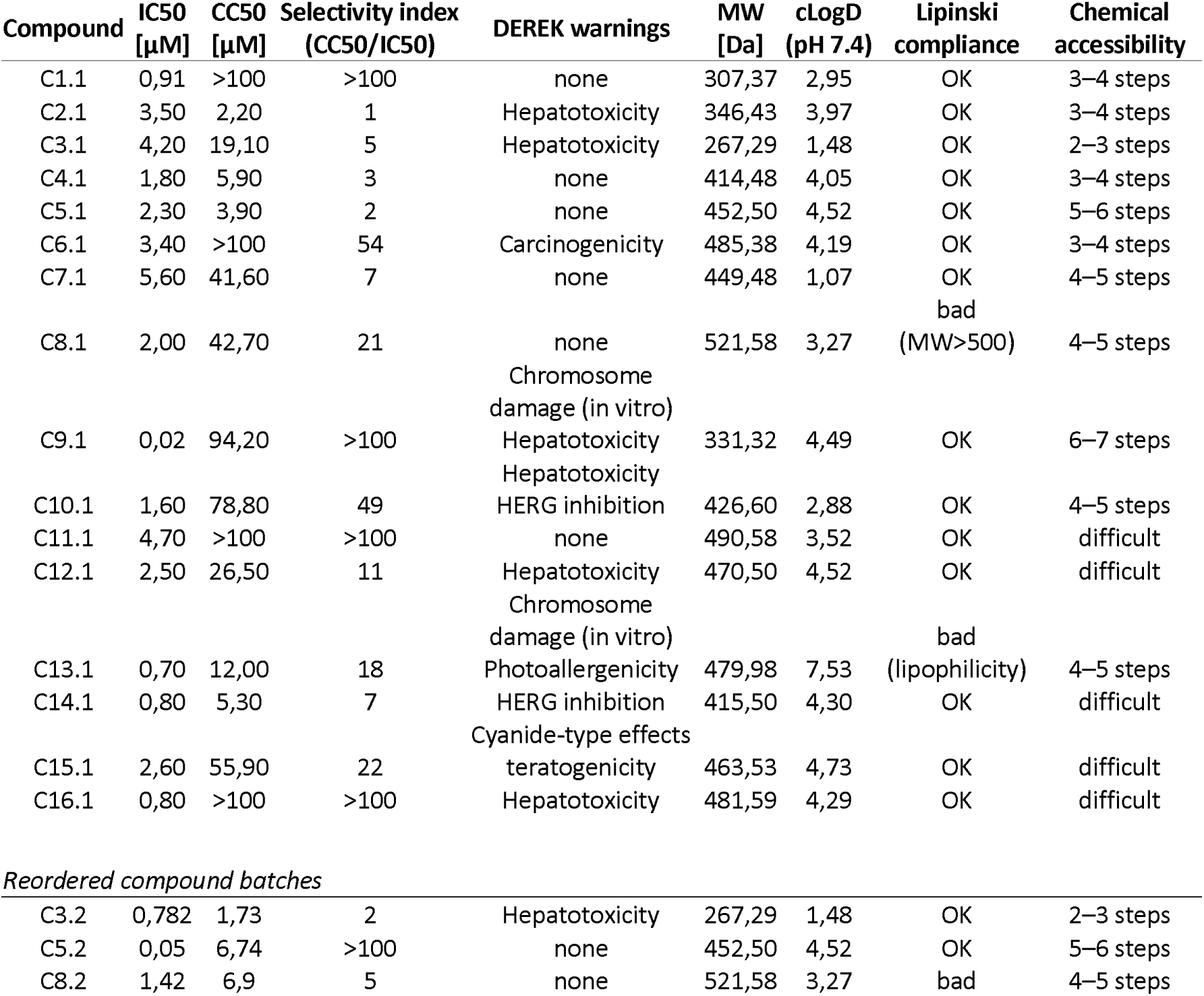
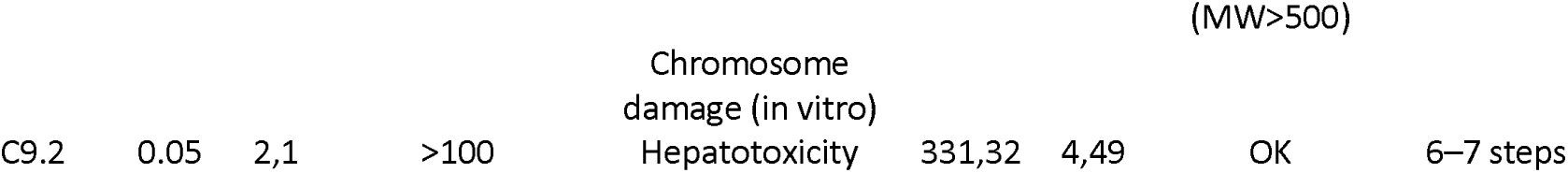
Pharmacological, predicted toxicity, physicochemical, and chemical accessibility profiles of BKPyV-active compounds.

### Integrated pharmacological and in silico profiling prioritizes compounds for further characterization

Because antiviral potency alone is insufficient for compound prioritization, the 16 inhibitors were further assessed according to predicted toxicity, physicochemical properties and synthetic accessibility. DEREK predictions were used to identify potential toxicity liabilities, while molecular weight and cLogD were considered together with Lipinski rule-of-five compliance and estimated synthetic accessibility (Table 1).

To support compound prioritization beyond antiviral potency alone, IC₅₀ and CC₅₀ values and the resulting selectivity indices were considered together with predicted toxicity liabilities, physicochemical properties, Lipinski compliance, and chemical accessibility (Table 1). The compounds displayed markedly different pharmacological profiles, with selectivity indices ranging from approximately 1 to >100. Eleven compounds (C1, C5-C6, C8-C13, C15, and C16) exhibited an SI >10 in at least one of the tested compound batches. However, high selectivity was not necessarily accompanied by an otherwise favorable compound profile. Several compounds with high SI values carried predicted toxicity liabilities, including hepatotoxicity, carcinogenicity, chromosome damage, HERG inhibition, photoallergenicity, or teratogenicity, whereas others were limited by high molecular weight, lipophilicity, or predicted chemical accessibility. Conversely, C1 and C11 combined SI values >100 with no DEREK toxicity warnings, although C11 was classified as difficult in terms of chemical accessibility. C8 showed an SI of 21 in the initial analysis but exceeded the Lipinski molecular-weight criterion (MW 521.58 Da), whereas C9 combined particularly high antiviral potency (IC₅₀ 0.02 µM) and an SI >100 with predicted chromosome-damage and hepatotoxicity warnings. These analyses identified compounds with distinct strengths and liabilities rather than a single uniformly superior candidate. C1 showed a particularly favorable pharmacological and predicted toxicity profile and therefore represents an attractive candidate for future optimization. For the mechanistic experiments described below, however, compounds were selected primarily to represent distinct functional phenotypes identified in the time-of-addition and pretreatment experiments rather than on the basis of developability alone.

### Selected inhibitors retain activity against another polyomavirus

To assess the breadth of antiviral activity and explore the potential for subsequent studies in murine models, compounds C1–C16 were additionally evaluated against the more distantly related murine polyomavirus (MuPyV). NMuMG cells were infected with MuPyV at an MOI of 0.5 and treated with the compounds at 10 µM concentration. Viral genome copies were quantified at 6 days post-infection in both culture supernatants and cell-associated fractions (Suppl. Fig. S3). Several compounds reduced MuPyV loads, demonstrating that the activity of a subset of compounds extends beyond BKPyV to a more distantly related polyomavirus. The differential activity observed across polyomaviruses further supports functional diversity within the compound panel and identifies compounds with broader anti-polyomavirus activity.

### Time-of-addition analysis distinguishes compounds according to the temporal window of antiviral activity

We next investigated how the timing of compound administration affected antiviral activity. All compounds were subjected to time-of-addition analysis in hTERT RPTE cells infected with BKPyV at an MOI of 0.1. Compounds were added at 0, 3, 6, 12, 24, 48, or 96 h post-infection, and extracellular viral DNA was quantified at 6 days post-infection. Based on the latest time point at which compound addition still resulted in a significant reduction in BKPyV load, the compounds could be grouped into three temporal activity classes. C16, C3, and C9 were selected as representative compounds displaying distinct time-of-addition profiles (Fig. 3A). C16 showed the strongest dependence on earlier administration, whereas C3 retained antiviral activity when added at later stages of infection. C9 displayed the latest effective treatment window, significantly reducing BKPyV load even when first added at 96 h post-infection. Thus, the compounds differed substantially in the time after infection at which treatment could still suppress progeny virus production, suggesting that their antiviral effects may involve different stages of the BKPyV replication cycle.

**Figure 3.**
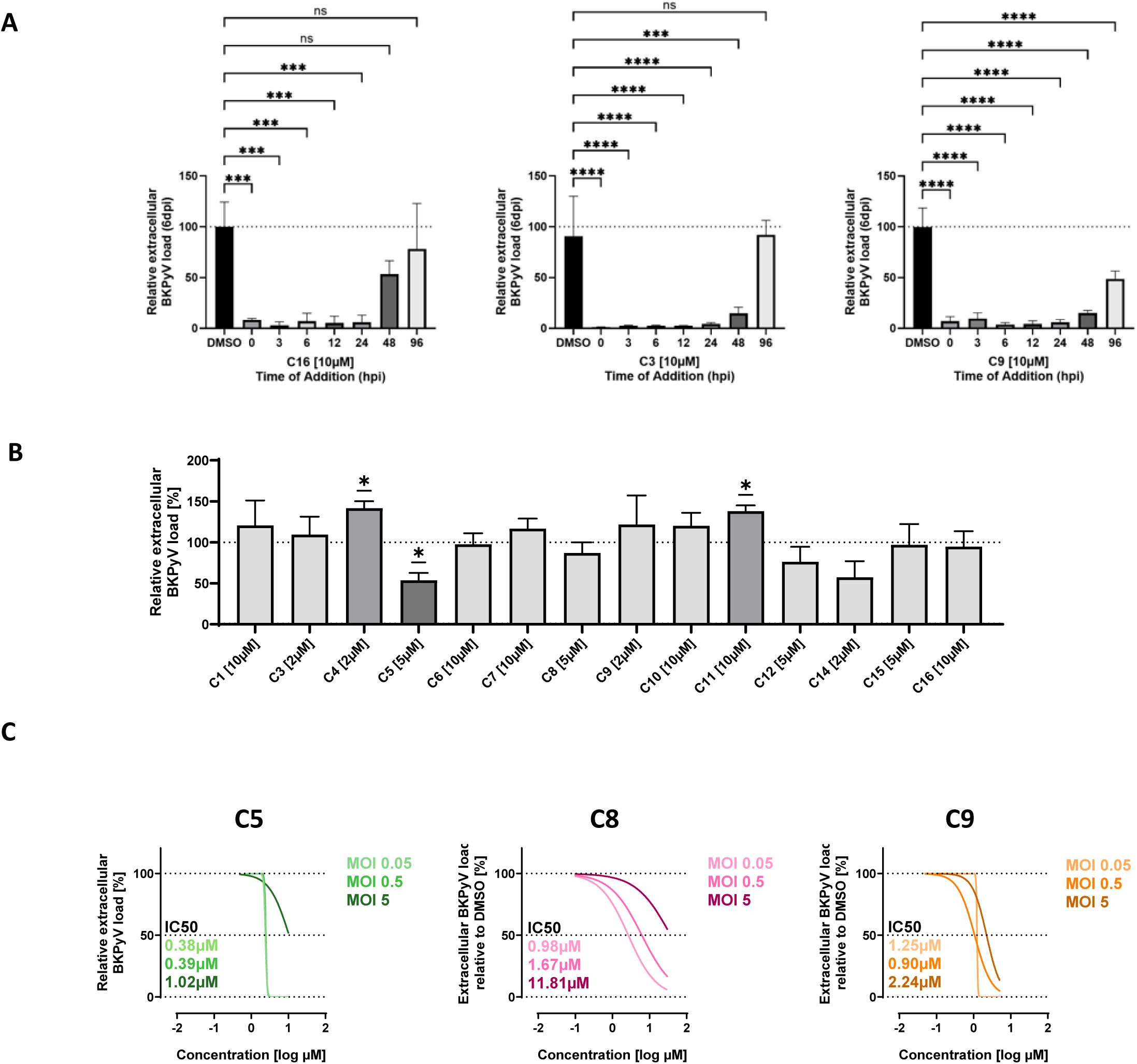
Temporal and MOI-dependent characterization of the antiviral activity of selected compounds against BKPyV. **(A)** Time-of-addition analysis of C16, C3, and C9. hTERT RPTE cells were infected with BKPyV (MOI 0.1) and treated with the indicated compounds (10 µM) at different time points post-infection (0, 3, 6, 12, 24, 48, and 96 hpi). Extracellular BKPyV loads were quantified by VP1 qPCR from cell culture supernatants at 6 days post-infection (dpi) and normalized to the DMSO-treated control (100%). Statistical significance was determined by one-way ANOVA followed by Dunnett’s multiple comparisons test versus DMSO. Bars represent mean ± SEM of biological replicates. ns, not significant; ***p < 0.001; ****p < 0.0001. **(B)** Pretreatment analysis of the indicated compounds. hTERT RPTE cells were pretreated with the respective compounds for 3 h, after which the compounds were removed by washing prior to BKPyV infection. Compound concentrations were selected based on their dose–response profiles and cytotoxicity values. Extracellular BKPyV release was assessed at 6 dpi by VP1 qPCR of cell culture supernatants. **(C)** MOI-dependent dose–response analysis of C5, C9, and C8. hTERT RPTE cells were infected with BKPyV at MOIs of 0.05, 0.5, and 5 and treated with increasing concentrations of the respective compound. Extracellular BKPyV loads were quantified by VP1 qPCR from cell culture supernatants at 6 dpi and normalized to the DMSO-treated control. Dose–response curves were fitted for each MOI, and the corresponding IC₅₀ values were calculated as indicated.

We additionally tested whether transient compound exposure before infection was sufficient to affect subsequent BKPyV replication. Cells were pretreated with the indicated compounds for 3 h at concentrations selected based on their individual dose–response and cytotoxicity profiles. Compounds were subsequently removed by washing prior to BKPyV infection, and extracellular viral DNA was quantified at 6 days post-infection (Fig. 3B). Among the compounds tested, only pretreatment with C5 resulted in a significant reduction in extracellular BKPyV load compared with the DMSO-treated control. Pretreatment with the remaining compounds did not reduce viral loads, whereas C4 and C11 resulted in a significant increase in extracellular BKPyV DNA. These findings indicate that, with the exception of C5, transient compound exposure prior to infection was not sufficient to reproduce the antiviral activity observed when compounds were present during infection.

Based on the time-of-addition and pretreatment analyses, C5, C8, and C9 were selected for subsequent mechanistic characterization because they represented distinct functional antiviral phenotypes. C5 was selected because it was the only compound for which transient pretreatment alone reduced subsequent BKPyV replication. C8 represented compounds whose antiviral activity was retained when treatment was initiated during the first 48 h of infection, whereas C9 represented the distinct late-acting phenotype, remaining effective even when treatment was initiated at 96 h post-infection. Selection of these compounds was therefore intended to capture mechanistically informative differences in antiviral behavior rather than to identify the three compounds with the most favorable overall drug-development profiles.

### C5, C8, and C9 display distinct relationships between antiviral potency and viral inoculum

To investigate whether antiviral potency was influenced by the amount of input virus, concentration– response analyses were performed for C5, C8, and C9 at BKPyV MOIs of 0.05, 0.5, and 5. Extracellular BKPyV DNA was quantified at 6 days post-infection, and IC₅₀ values were determined independently for each MOI (Fig. 3C).

C5 displayed a steep concentration–response profile across all three infection conditions. The fitted IC₅₀ values increased from 0.35 µM at an MOI of 0.05 to 0.59 µM at an MOI of 0.5 and 1.64 µM at an MOI of 0.5. Because of the unusually steep shape of the C5 concentration–response curves, however, the magnitude of this apparent MOI-dependent shift should be interpreted cautiously. In contrast, C8 showed comparatively little variation in antiviral potency across the tested viral inocula, with IC₅₀ values of 0.96, 1.67, and 1.61 µM at MOIs of 0.05, 0.5, and 5, respectively. C9 showed a more gradual increase in IC₅₀ with increasing viral input, from 1.25 µM at an MOI of 0.05 to 1.90 µM at an MOI of 0.5 and 2.21 µM at an MOI of 5.

Overall, the three compounds displayed distinct relationships between viral inoculum and fitted antiviral potency. C8 remained comparatively stable across increasing MOIs, whereas C9 showed a modest inoculum-dependent shift. Although the fitted IC₅₀ values for C5 increased with viral input, the steep concentration–response profile limits conclusions regarding the magnitude of this effect. Overall, the three compounds displayed distinct relationships between viral inoculum and fitted antiviral potency. C8 remained comparatively stable across increasing MOIs, whereas C9 showed a modest inoculum-dependent shift. Although the fitted IC₅₀ values for C5 increased with viral input, the steep concentration–response profile limits conclusions regarding the magnitude of this effect. Changes in apparent potency with viral inoculum alone cannot distinguish between viral and cellular targets. However, considered together with the C5 pretreatment phenotype and the compound-induced host transcriptional responses, these findings are compatible with mechanisms involving host-cell processes.

### C5, C8 and C9 exert distinct effects on BKPyV attachment and viral gene expression

To determine whether the selected inhibitors affect an early step of BKPyV infection, we examined the attachment of fluorescently labeled BKPyV VLPs to hTERT RPTE cells in the presence of C5, C8, or C9. Cells were incubated with 7*10^5 transducing units fluorescently labeled VLPs per cell (MOI 1) in the presence of C5 (5µM), C8 (5µM), or C9 (2µM), or the corresponding DMSO control. Following incubation for 2hrs at 4°C, cell-associated VLP fluorescence was quantified by flow cytometry (Fig. 4A–B). C5 and C8 resulted in VLP-associated fluorescence distributions comparable to the DMSO control, indicating no detectable reduction in VLP attachment under these conditions. In contrast, C9 markedly increased cell-associated VLP fluorescence, resulting in a higher mean fluorescence intensity compared with DMSO-treated cells. Thus, none of the three compounds reduced BKPyV VLP attachment, whereas C9 unexpectedly increased the amount of cell-associated VLP signal.

**Figure 4:**
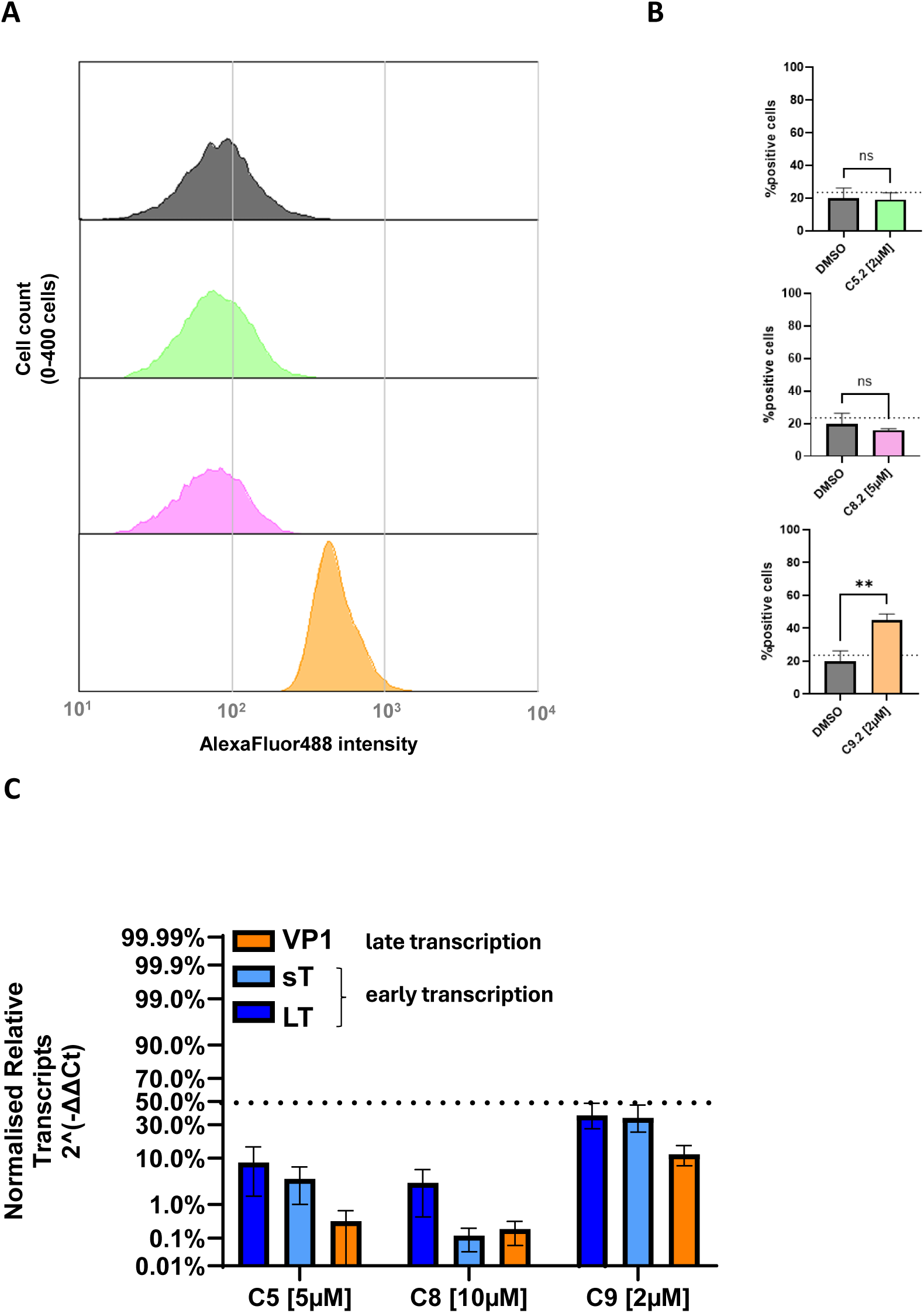
Effects of selected antiviral compounds on BKPyV VLP attachment and viral gene expression. **(A)** Representative flow cytometry histograms of BK polyomavirus (BKPyV) virus-like particle (VLP) attachment to hTERT RPTE cells in the presence of DMSO or the indicated compounds. Cells were incubated with fluorescently labeled BKPyV VLPs in the presence of DMSO (gray), C5 (green), C8 (pink), or C9 (orange), and cell-associated VLP fluorescence was measured by flow cytometry. Histograms show the distribution of VLP-associated fluorescence (APC) as cell count normalized to the maximum event count. **(B)** Quantification of BKPyV VLP attachment shown in (A), expressed as the mean fluorescence intensity (MFI) of cell-associated VLPs for DMSO-, C5-, C8-, and C9-treated cells. Bars represent mean ± SEM. **(C)** Effect of C5, C8, and C9 on BKPyV viral transcript levels. hTERT RPTE cells were infected with BKPyV and treated with the indicated compounds. At 24 h post-infection, RNA was isolated and transcript levels of the early viral genes encoding large T antigen (LT) and small t antigen (sT), as well as the late capsid protein VP1, were quantified by RT–qPCR. Transcript levels were normalized to the DMSO-treated control (set to 100%); the dotted line indicates 50% of the control level. Bars represent mean ± SEM.

Having examined the effects of C5, C8, and C9 on VLP attachment, we next investigated whether the compounds affected viral gene expression during the early phase of BKPyV infection. hTERT RPTE cells were infected with BKPyV and treated with C5, C8, or C9 at the time of infection. At 24 h post-infection, transcript levels of the early viral genes LT and sT and the late capsid gene VP1 were quantified by RT–qPCR and normalized to DMSO-treated infected cells (Fig. 4C).

The three compounds displayed distinct effects on early (LT and sT) and late (VP1) viral transcript abundance. C8 reduced both early and late viral transcripts, with the reduction in VP1 being more pronounced than that observed for the early transcripts. C5 similarly reduced viral transcript abundance, with a pronounced effect on the late VP1 transcript in addition to a reduction in early transcript levels. In contrast, C9 had little effect on early LT and sT transcript abundance, whereas the late VP1 transcript was moderately reduced. Thus, C5 and C8 affected both early and late viral gene expression, whereas the transcriptional phenotype of C9 was predominantly associated with the late viral transcript.

Together with the VLP attachment experiments, these results demonstrate distinct effects of C5, C8, and C9 during early BKPyV infection and indicate that their antiviral activity is not attributable to inhibition of initial virus attachment. Instead, the differential effects on early and late viral transcript abundance suggest that the compounds interfere with distinct processes occurring downstream of virus attachment.

### BKPyV infection induces a distinct host transcriptional program in hTERT RPTE cells

Having established distinct effects of C5, C8, and C9 on viral infection phenotypes, we next examined how BKPyV infection and compound treatment affected the host-cell transcriptional state. As a reference for subsequent comparison with the compound-associated responses, we first defined the transcriptional signature of BKPyV infection in hTERT RPTE cells. Transcriptomic profiling was therefore performed in BKPyV-infected and non-infected cells at 48 h post-infection, with both conditions maintained in the presence of DMSO. Principal component analysis (PCA) revealed a clear separation between BKPyV-infected and non-infected samples, with PC1 accounting for 61.5% and PC2 for 20.3% of the total variance (Suppl. Fig. S4A). Despite some variation between the two biological replicates, both infected samples were clearly separated from the non-infected controls along PC1, indicating that BKPyV infection was the major source of transcriptional variation in the dataset.

Differential expression analysis further demonstrated a substantial host transcriptional response to BKPyV infection. At a false discovery rate (FDR) of 0.05, 774 genes were differentially expressed, comprising 473 upregulated and 301 downregulated genes in BKPyV-infected compared with non-infected DMSO-treated cells (Suppl.Fig. S4B). These differentially expressed genes were subsequently used to define the BKPyV infection-associated transcriptional signature and to investigate the cellular processes associated with infection (Fig. 5).

**Figure 5.**
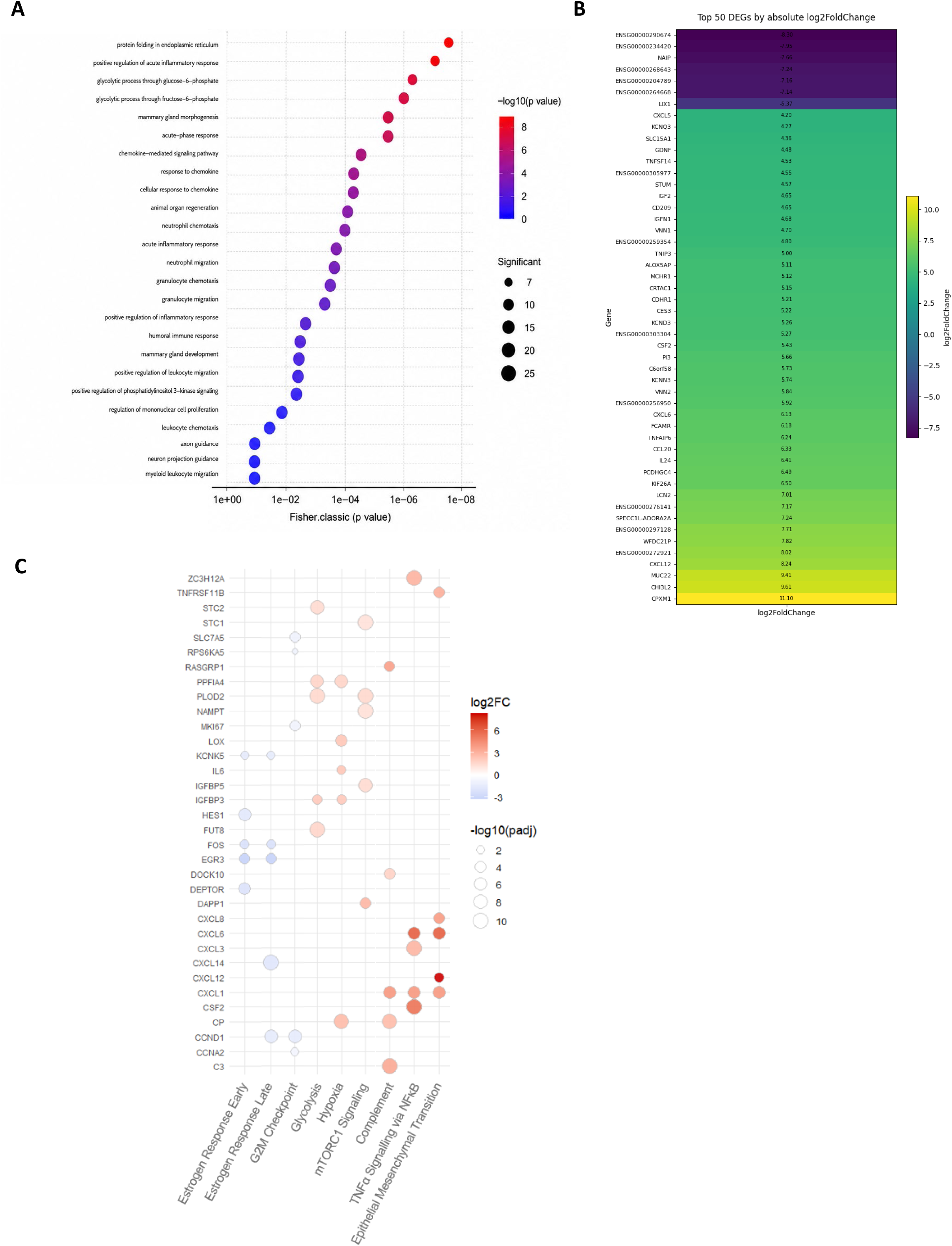
Transcriptional signature and pathway alterations associated with BKPyV infection in hTERT RPTE cells. Transcriptomic profiling was performed in hTERT RPTE cells 48 h post-infection by comparing BKPyV-infected with non-infected control cells, both maintained in the presence of DMSO. **(A)** Gene Ontology (GO) Biological Process enrichment analysis of differentially expressed genes (DEGs) associated with BKPyV infection. Enriched biological processes are ranked according to Fisher’s exact p value. Dot color represents −log10(p value), and dot size indicates the number of significant genes associated with each term. **(B)** Heatmap showing the 50 DEGs with the largest absolute log2 fold changes between BKPyV-infected and non-infected cells. Color represents the direction and magnitude of differential expression, with red and blue indicating increased and decreased expression, respectively, in BKPyV-infected relative to non-infected cells. **(C)** Gene set enrichment analysis (GSEA) of the BKPyV infection-associated transcriptional signature using the Hallmark gene set collection. Pathways meeting the enrichment criteria (adjusted p < 0.1, |NES| > 1.5, and gene set size ≥10) are shown together with the five leading-edge genes exhibiting the largest absolute log2 fold changes within each pathway. Rows represent individual genes and columns represent enriched Hallmark pathways. Dot color indicates the direction and magnitude of differential expression (log2 fold change), with red and blue representing increased and decreased expression, respectively, in BKPyV-infected relative to non-infected cells. Dot size represents −log10(adjusted p value).

Gene Ontology (GO) Biological Process analysis revealed significant enrichment of several cellular processes within the BKPyV infection-associated signature (Fig. 5A). Among the most significantly enriched terms were protein folding in the endoplasmic reticulum, glycolytic processes, and positive regulation of acute inflammatory responses. A substantial proportion of the enriched terms was related to inflammatory and chemokine-associated responses, including chemokine-mediated signaling, response to chemokines, cellular response to chemokines, acute inflammatory response, and positive regulation of inflammatory responses. Processes associated with immune-cell migration, including neutrophil chemotaxis and migration, granulocyte chemotaxis and migration, leukocyte chemotaxis, and myeloid leukocyte migration, were also enriched. In addition, the infection-associated signature included processes related to phosphatidylinositol 3-kinase signaling and cellular metabolic responses.

Consistent with the broad transcriptional response identified by the enrichment analysis, examination of the 50 DEGs with the largest absolute log2 fold changes revealed pronounced changes in individual host transcripts (Fig. 5B). Both strongly increased and decreased transcripts were present, although upregulated genes predominated among the genes displaying the largest positive expression changes. Several genes associated with inflammatory and chemokine responses were among the strongly induced transcripts, consistent with the functional enrichment observed at the pathway level.

To further resolve the genes contributing to these infection-associated processes, differentially expressed genes were mapped to selected enriched functional categories (Fig. 5C). Multiple chemokine- and inflammation-associated genes, including CXCL1, CXCL6, CXCL8, and CXCL12, showed increased expression and contributed to several of the enriched categories. Additional differentially expressed genes, including PLOD2, LOX, IGFBP3, FSTL1, and PTX3, were represented across processes associated with cellular stress and tissue or extracellular organization.

This signature in BKPyV infected RPTE cells was subsequently used as a reference to determine how treatment with the prioritized antiviral compounds modifies cellular transcriptional responses in non-infected and BKPyV-infected cells.

### C9 induces a distinct host transcriptional response that intersects with the BKPyV infection signature

Transcriptomic profiling of C5-, C8-, and C9-treated hTERT RPTE cells revealed compound-specific alterations in host gene expression (Fig 6, suppl. Fig. S5). In conjunction with the preceding functional analyses, C9 displayed a particularly distinct phenotype. While C5 and C8 showed antiviral activity predominantly when treatment was initiated during earlier phases of infection, C9 retained activity when added at comparatively late time points. Moreover, C9, but not C5 or C8, markedly altered cell-associated BKPyV VLP fluorescence, further distinguishing its phenotype during early virus–cell interactions. We therefore examined the C9-associated host response in greater detail and assessed its relationship to the BKPyV infection-associated transcriptional signature.

**Figure 6.**
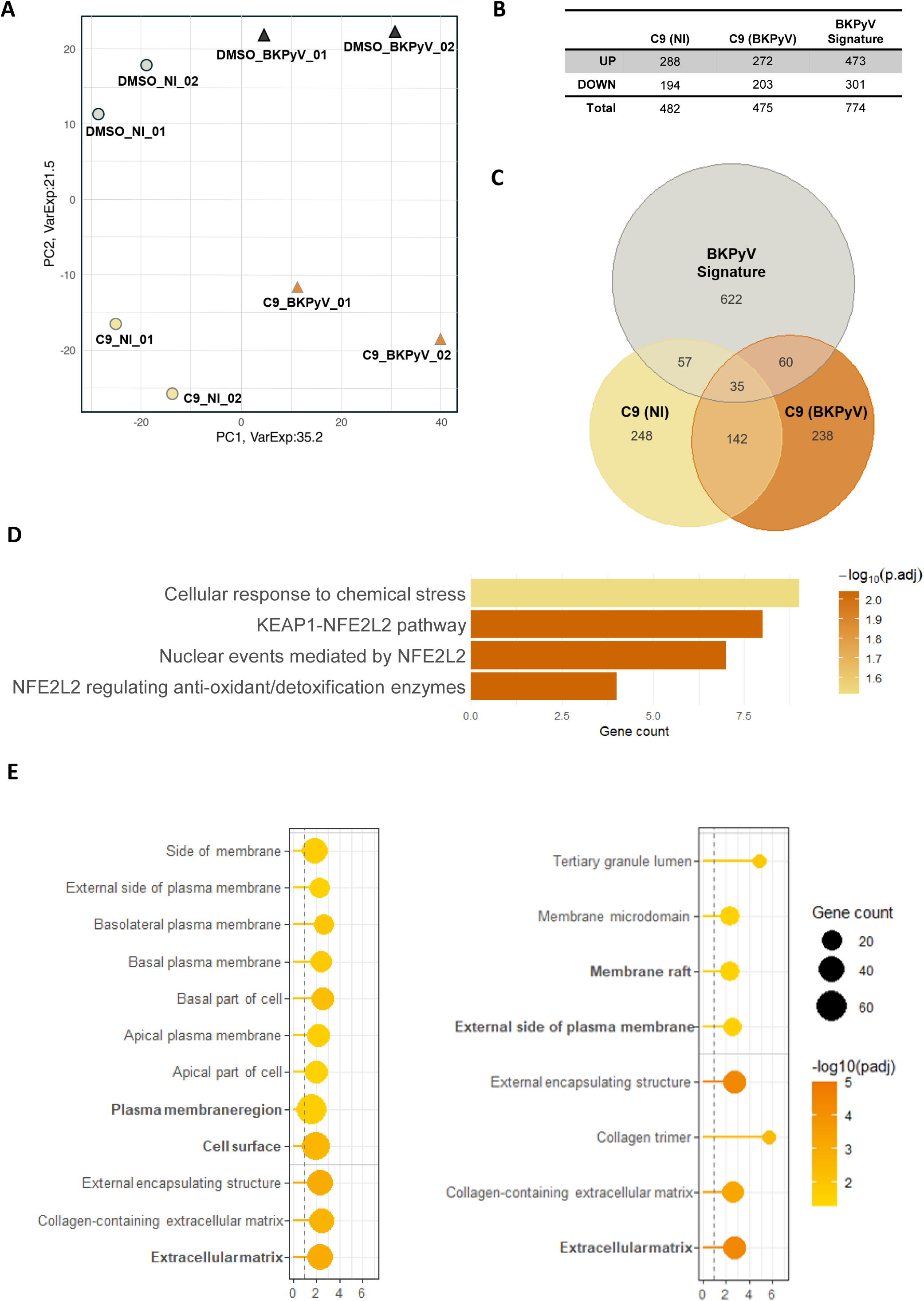
Transcriptomic characterization of the cellular response to C9 treatment in non-infected and BKPyV-infected hTERT RPTE cells. Transcriptomic profiling was performed in hTERT RPTE cells 48 h post-infection to characterize C9-induced changes in non-infected (NI) and BKPyV-infected cells. **(A)** Principal component analysis (PCA) of DMSO- and C9-treated non-infected and BKPyV-infected cells. Each point represents an individual biological replicate. PC1 and PC2 account for 35.2% and 21.5% of the total variance, respectively. **(B)** Number of differentially expressed genes (DEGs) identified for the indicated transcriptional signatures. **(C)** Venn diagram showing the overlap between the BKPyV infection-associated transcriptional signature and the transcriptional responses to C9 treatment in non-infected [C9 (NI)] and BKPyV-infected [C9 (BKPyV)] cells. Numbers indicate genes unique to or shared between the respective signatures. **(D)** Functional enrichment analysis of genes associated with the C9-induced transcriptional response, showing significantly enriched pathways related to cellular responses to chemical stress and NRF2-associated signaling. Bars indicate gene counts, and color represents −log10(adjusted p value). **(E)** Gene Ontology Cellular Component enrichment analysis of the C9-induced transcriptional signatures in non-infected (left) and BKPyV-infected (right) cells. Enriched cellular components are plotted according to fold enrichment. Dot size represents the number of genes associated with each term, and dot color indicates −log10(adjusted p value).

Principal component analysis showed a pronounced separation of C9-treated samples from the corresponding DMSO-treated controls in both non-infected and BKPyV-infected cells (Fig. 6A). C9 treatment therefore induced substantial changes in the cellular transcriptional state irrespective of infection status. Comparison of the resulting differentially expressed gene sets with the BKPyV infection signature revealed both distinct and overlapping transcriptional responses (Fig. 6B–C). Of the differentially expressed genes, 248 were unique to C9-treated non-infected cells and 238 were unique to C9-treated BKPyV-infected cells, whereas 622 were unique to the BKPyV infection signature. Importantly, 142 genes were shared between the C9 responses in non-infected and BKPyV-infected cells, consistent with a substantial infection-independent component of the C9-induced transcriptional response. In addition, 57 genes were shared between the C9 response in non-infected cells and the BKPyV infection signature, 60 between the C9 response in infected cells and the BKPyV infection signature, and 35 genes were common to all three signatures.

Functional enrichment analysis further indicated that the C9-induced transcriptional response involved cellular stress-associated processes (Fig. 6D). Enriched terms included *cellular response to chemical stress*, the *KEAP1–NFE2L2 pathway, nuclear events mediated by NFE2L2, and NFE2L2-regulated antioxidant/detoxification enzymes*. Thus, C9 treatment was associated with a transcriptional program involving the KEAP1–NFE2L2/NRF2 stress-response axis, although these data do not establish this pathway as the direct molecular target of C9.

Gene Ontology Cellular Component analysis additionally identified enrichment of membrane- and extracellular matrix-associated categories following C9 treatment (Fig. 6E). In non-infected cells, enriched terms included the plasma membrane region, cell surface, external side of the plasma membrane, and extracellular matrix. Related categories were also enriched in C9-treated BKPyV-infected cells, including membrane rafts, the external side of the plasma membrane, collagen-containing extracellular matrix, and extracellular matrix.

Together, these data show that C9 induces a pronounced host transcriptional response that is largely retained in the absence of BKPyV infection and partially intersects with cellular programs altered during infection. The infection-independent nature of a substantial component of this response, together with the distinct phenotype of C9 observed in the preceding functional assays, further supports a predominantly host-associated mechanism of antiviral activity and prompted a more detailed comparison of the C9-induced and BKPyV infection-associated transcriptional programs.

### C9 counteracts BKPyV-associated transcriptional changes in extracellular matrix and EMT-related programs

Given the partial intersection between the C9-induced and BKPyV infection-associated transcriptional responses, we next examined whether C9 affected specific cellular programs altered during BKPyV infection. Comparative analysis revealed a strikingly opposing pattern within EMT-associated transcriptional programs (Fig. 7A). BKPyV infection was predominantly associated with increased expression of EMT-related genes, whereas C9 treatment of non-infected cells was associated with decreased expression of a partially overlapping set of EMT-associated genes. BKPyV infection was associated predominantly with increased expression of EMT-related genes, including SAT1, PLOD2, NID2, LOX, IGFBP3, TNFRSF11B, PTX3, CXCL1, CXCL6, CXCL8, and CXCL12. In contrast, several EMT-associated genes within the C9 response in non-infected cells, including SFRP1, ABI3BP, TGFBI, TAGLN, PLOD2, COL4A1, COL11A1, DAB2, PVR, VCAN, and FSTL1, showed decreased expression. Thus, although both BKPyV infection and C9 treatment affected genes associated with EMT, the transcriptional responses were largely distinct and frequently differed in direction.

**Fig 7.**
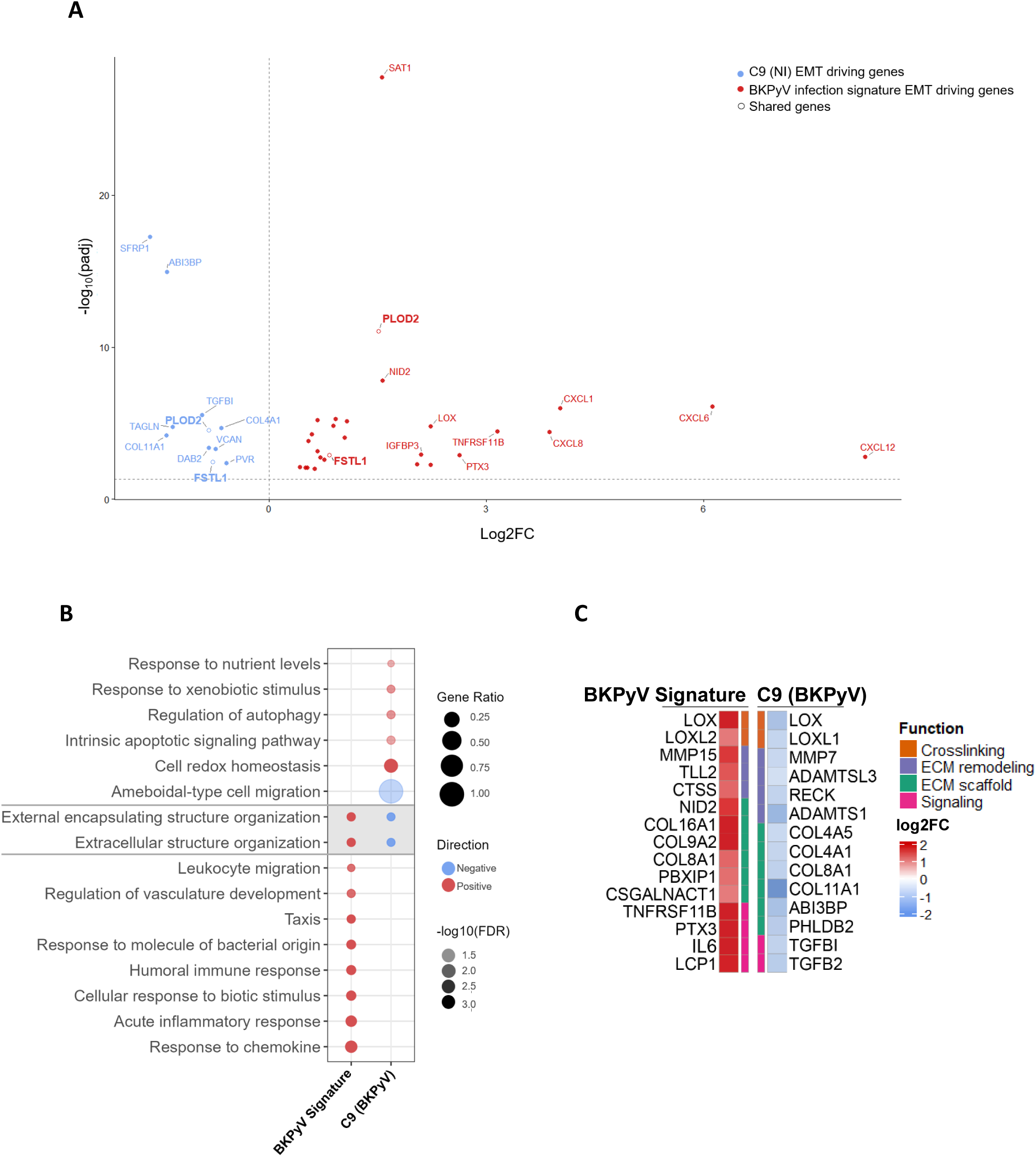

Functional enrichment analysis further supported this divergence (Fig. 7B). BKPyV infection was positively associated with multiple inflammatory and immune-response processes, including leukocyte migration, response to molecule of bacterial origin, humoral immune response, cellular response to biotic stimulus, acute inflammatory response, and response to chemokine. In contrast, C9 treatment in non-infected cells was associated with negative regulation of processes related to ameboid-type cell migration, external encapsulating structure organization, and extracellular structure organization. C9 treatment was additionally associated with changes in cellular processes including response to nutrient levels, response to xenobiotic stimulus, regulation of autophagy, intrinsic apoptotic signaling, and cellular redox homeostasis.

A focused comparison of genes associated with extracellular matrix organization and related functions further demonstrated opposing transcriptional patterns between BKPyV infection and C9 treatment (Fig. 7C). Genes classified into functional groups related to ECM scaffold, ECM remodeling, crosslinking, and signaling were predominantly increased within the BKPyV infection-associated signature, whereas the corresponding genes showed reduced expression following C9 treatment in non-infected cells. This inverse pattern indicates that C9 does not globally reverse the BKPyV infection signature but instead counteracts a defined subset of infection-associated transcriptional changes, particularly those related to extracellular matrix organization and remodeling.

Together, these analyses identify ECM/EMT-associated transcriptional remodeling as a point of convergence between BKPyV infection and the cellular response to C9, but with predominantly opposing regulatory effects. Combined with the infection-independent transcriptional response to C9 and its effects observed in the preceding functional assays, these findings support a model in which C9 alters host-cell programs that are also modulated during BKPyV infection, rather than directly reproducing or broadly suppressing the infection-associated transcriptional response.

## Discussion

Despite the substantial clinical burden of BKPyV in immunocompromised patients, treatment remains largely limited to reduction of immunosuppression, and no specific antiviral therapy is currently available. Although emerging approaches, including broadly neutralizing antibodies (14, 15), offer new therapeutic perspectives, effective pharmacological options remain limited. In this study, we used an unbiased phenotypic screening strategy to identify and characterize small-molecule inhibitors of polyomavirus infection. The identified compounds displayed distinct pharmacological and temporal profiles, supporting the potential of targeting host-dependent processes to restrict BKPyV replication.

The use of SV40 as a surrogate enabled phenotypic compound discovery without requiring a predefined viral or cellular target. Previous polyomavirus screening approaches have included target-based strategies, such as the identification of bithionol and hexachlorophene as inhibitors of large T antigen ATPase activity and subsequent SV40 and BKPyV replication (31). In our study, approximately half of the SV40-active compounds retained antiviral activity against BKPyV in renal epithelial cells, supporting the utility of SV40 for initial compound discovery while also emphasizing the importance of subsequent validation in a BKPyV-relevant cellular system.

The BKPyV-active compounds did not, however, behave as a homogeneous class of inhibitors. This functional diversity is particularly relevant in the context of polyomavirus biology, as the limited coding capacity of BKPyV results in extensive dependence on cellular machinery throughout its replication cycle, including entry and intracellular trafficking, capsid disassembly, nuclear transport, transcription, DNA replication, and progeny production. Phenotypic screening may therefore capture inhibitors of cellular processes required at different stages of infection that would not readily emerge from approaches directed toward individual viral proteins. Consistent with this concept, the identified compounds differed in their activity across polyomaviruses, temporal window of efficacy, response to pretreatment, dependence of antiviral potency on viral inoculum, and effects on early virus–cell interactions and viral transcription. Together, these distinct pharmacological phenotypes suggested that the compounds interfere with different processes required for productive BKPyV infection. The unique pretreatment phenotype of C5 provided further evidence for a cellular contribution to antiviral activity, as transient exposure before infection was sufficient to reduce subsequent BKPyV replication, consistent with induction of a less permissive cellular state. C5, C8, and C9 also retained concentration-dependent antiviral activity across a 100-fold range of viral input, although their IC₅₀ values varied with MOI. While these pharmacological observations do not independently distinguish cellular from viral targets, together with the compound-induced host transcriptional responses they favor mechanisms involving cellular processes rather than exclusive inhibition of a viral target. Direct target-identification studies will be required to define the underlying molecular mechanisms. The experiments with fluorescent BKPyV VLPs further differentiated the prioritized compounds. C5 and C8 did not substantially alter cell-associated VLP fluorescence, making inhibition of initial particle attachment unlikely to account for their antiviral activity under the conditions tested. C9, in contrast, markedly increased the cell-associated VLP signal. This phenotype is particularly interesting because it argues against simple competitive inhibition of viral receptor binding. BKPyV entry depends on interactions with sialylated gangliosides and subsequent intracellular trafficking toward the endoplasmic reticulum, and therefore alterations in membrane composition, particle internalization, retention, or trafficking could change the amount of cell-associated VLP fluorescence without increasing productive infection. The present assay cannot discriminate between these possibilities. Accordingly, the C9 phenotype should be interpreted as an alteration of early virus–cell interactions rather than evidence for enhanced viral attachment per se. The broader cellular consequences of BKPyV infection and compound treatment became apparent from transcriptomic profiling. BKPyV infection generated a distinct transcriptional signature in hTERT RPTE cells, including changes in inflammatory and chemokine-associated processes, metabolism, cellular stress responses, and extracellular organization. Notably, despite this broad response, we did not observe a prominent canonical interferon signature. This is consistent with previous studies showing limited induction of interferon-stimulated genes during productive BKPyV infection of renal proximal tubular epithelial cells, suggesting that BKPyV can evade or attenuate innate antiviral signaling in its natural target cells (32–34). Previous transcriptomic studies have likewise demonstrated extensive remodeling of renal epithelial cells during BKPyV infection, including infection-dependent cellular heterogeneity and alterations in stress and mitochondrial programs (32, 34). The use of hTERT RPTE cells should nevertheless be considered when interpreting these signatures, as cultured primary and immortalized RPTE cells display features of stressed or injured renal epithelium rather than fully differentiated proximal tubular cells *in vivo*, although comparable BKPyV-induced responses have been reported in primary and immortalized RPTE cells (35). Together, our findings support substantial but selective remodeling of renal epithelial cell physiology during BKPyV infection, characterized by inflammatory, metabolic, and stress-associated responses without a correspondingly strong canonical interferon signature.

The prominent inflammatory and chemokine-associated component of the BKPyV signature may also be relevant to the response observed in BKPyV-associated nephropathy *in vivo*. Transcriptomic analyses of affected renal allografts have demonstrated strong inflammatory gene expression, including cytokine- and chemokine-associated pathways, with substantial overlap with transcriptional patterns observed during acute cellular rejection (36, 37). Thus, despite the limitations of cultured renal epithelial models, the inflammatory signature observed in our study recapitulates features reported in BKPyV-infected renal tissue and suggests that infected tubular epithelial cells may contribute directly to the inflammatory environment associated with BKPyV nephropathy.

Of particular interest was the association of BKPyV infection with extracellular matrix (ECM)- and epithelial–mesenchymal transition (EMT)-related transcriptional programs. Similar changes have been reported in BKPyV nephropathy, including increased expression of matrix collagens, TGF-β-associated pathways, matrix metalloproteinases, and EMT-associated markers in affected renal allografts (37, 38). Our findings suggest that transcriptional remodeling of infected renal epithelial cells may contribute to these tissue-level changes, although the fibrotic response in vivo necessarily reflects interactions between epithelial, immune, and stromal compartments.

Transcriptomic profiling of C5, C8, and C9 revealed distinct cellular responses, consistent with their different antiviral phenotypes. These signatures provide hypotheses regarding cellular processes associated with compound activity rather than evidence for direct molecular targets, particularly because transcriptional changes measured 48 h after treatment may include both primary and secondary responses. Importantly, the three compounds should not be interpreted as equivalent candidates for further drug development. C5 combined a distinct pretreatment phenotype with antiviral activity in the absence of a comparably extensive transcriptional perturbation, whereas C8 likewise produced a more restricted host transcriptional response. C9 was selected for deeper pathway-level analysis because it generated the most distinctive mechanistic phenotype, not because it represented the most favorable development candidate. Indeed, its predicted toxicological liabilities and pronounced cellular stress response argue for particular caution in interpreting C9 as a therapeutic lead. C9 was particularly informative as a mechanistic probe because its transcriptional phenotype converged with several observations from the functional assays. C9 retained antiviral activity following comparatively late addition and markedly altered cell-associated BKPyV VLP fluorescence. In addition, a substantial component of the C9-induced transcriptional response was shared between infected and non-infected cells, indicating that these changes were not simply secondary to inhibition of viral replication. Together, these observations provided the rationale for examining in greater detail how C9-induced cellular changes intersect with the BKPyV-associated transcriptional program.

C9 treatment was associated with enrichment of cellular stress responses, including the KEAP1– NFE2L2/NRF2 axis. As NRF2 represents a broad adaptive response to chemical and metabolic stress, this enrichment does not identify NRF2 or KEAP1 as a molecular target of C9. Rather, it indicates that cellular stress adaptation forms part of the C9-induced response. This interpretation is particularly relevant given the predicted toxicological liabilities of C9 and therefore warrants caution in distinguishing mechanism-associated cellular responses from more general compound-induced stress. A more intriguing relationship emerged from the comparison of C9-induced and BKPyV-associated ECM/EMT programs. BKPyV infection was associated with increased expression of genes involved in extracellular matrix organization, remodeling, and EMT-associated processes, whereas C9 treatment produced predominantly opposing changes within a subset of these programs. Given that profibrotic and EMT-associated transcriptional changes have also been reported in BKPyV nephropathy (37), C9 appears to counteract a defined component of the BKPyV-associated cellular response rather than globally reversing the infection signature.

These transcriptional changes may also relate to the altered VLP phenotype observed with C9. Modulation of the extracellular matrix, plasma membrane, or cell-surface composition could influence BKPyV–cell interactions and may contribute to the increased cell-associated VLP signal.

However, because C9 remained antiviral when added comparatively late after infection, altered particle attachment alone is unlikely to explain its activity. C9 may therefore perturb a cellular process relevant at multiple stages of infection, although the VLP and transcriptional phenotypes could also represent independent consequences of a broader cellular perturbation. Distinguishing these possibilities will require direct analysis of particle internalization and trafficking together with functional validation of the pathways identified by transcriptomic profiling.

The activity of several compounds against the more distantly related MuPyV suggests that their antiviral effects may extend across polyomaviruses and could involve cellular requirements shared between these viruses. Beyond its mechanistic relevance, activity against MuPyV provides an opportunity for subsequent *in vivo* evaluation, as murine models of productive intrarenal polyomavirus infection and nephropathy are available (39). Compounds active against both BKPyV and MuPyV may therefore be particularly suitable for further preclinical characterization.

Several limitations should nevertheless be considered. Most importantly, the direct molecular targets of the identified compounds remain unknown. Evidence for host-associated mechanisms is based on converging pharmacological and transcriptional observations rather than direct target engagement, and pathway enrichment should therefore be considered hypothesis-generating. Genetic perturbation and orthogonal target-identification approaches will be required to establish the cellular factors responsible for antiviral activity. In addition, the present study was performed predominantly in cell culture. Although cytotoxicity measurements and selectivity indices provide an initial indication of the therapeutic window, they cannot predict systemic tolerability, pharmacokinetics, or renal exposure *in vivo*. Together with the predicted liabilities of some compounds and the batch-dependent differences observed for C5, C8, and C9, these considerations position the compounds primarily as starting points for mechanistic investigation and chemical optimization rather than as immediately developable antiviral candidates.

Our study identifies pharmacologically and functionally distinct inhibitors of BKPyV infection and highlights host-dependent processes as an underexplored source of antiviral targets. The compound panel has value both as a source of candidates for further optimization and as a set of chemical probes for dissecting BKPyV–host interactions. Compounds such as C1 and C5 combine favorable antiviral characteristics with comparatively limited predicted or transcriptional liabilities and therefore warrant further pharmacological refinement, whereas the pronounced cellular response elicited by C9 makes this compound particularly informative as a mechanistic probe. Defining the direct molecular targets responsible for these distinct antiviral phenotypes will be an important next step toward both mechanistic understanding and development of host-directed strategies against BKPyV.

## Material and Methods

### Cell culture

African green monkey kidney CV-1 cells (ATCC CCL-70) were used for establishment and execution of the primary phenotypic screening assay using simian virus 40 (SV40) as a surrogate polyomavirus. Primary human renal proximal tubular epithelial (RPTE) cells (Lonza, CC-2553) were used for secondary validation and pharmacological characterization of antiviral activity against BK polyomavirus (BKPyV). Immortalized human renal proximal tubular epithelial cells (RPTEC/TERT1; ATCC CRL-4031) were used for subsequent mechanistic and transcriptomic analyses. Normal murine mammary gland epithelial cells (NMuMG; ATCC CRL-1636) were used to assess antiviral activity against murine polyomavirus (MuPyV).

Cells were maintained at 37°C in a humidified atmosphere containing 5% CO₂. CV-1 and NMuMG cells were cultured in Dulbecco’s modified Eagle medium (DMEM) supplemented with 10% fetal bovine serum (FBS, Gibco) and 1% penicillin/streptomycin. RPTEC/TERT1 cells were maintained in a 1:1 mixture of DMEM and Ham’s F-12 (Sigma-Aldrich) supplemented with GlutaMAX/L-glutamine (1×), insulin-transferrin-selenium (1×) (Gibco), 1% penicillin/streptomycin (Gibco), 10 ng/mL epidermal growth factor (Gibco), 36 ng/mL hydrocortisone (Stemcell), 0.1 mg/mL G418 (Gibco), and 2% FBS. Primary RPTE cells were cultured in renal epithelial basal medium (REBM) (Gibco) supplemented with 2% FBS and gentamicin/amphotericin B (Gibco).

Cells were routinely passaged using 0.05% trypsin-EDTA (Gibco) and seeded at the density required for the respective experiment. BKPyV infections were generally initiated 16–24 h after seeding when cells had reached approximately 50–80% confluence.

### Viruses and determination of viral titers

SV40 (40) and BKPyV (Dunlop strain) stocks were propagated in CV-1 and WI-38 cells, respectively. Following propagation, virus-containing material was subjected to three freeze-thaw cycles, clarified by centrifugation, passed through a 0.22-µm filter, aliquoted, and stored at −80°C. MuPyV stocks were propagated in NMuMG cells and stored at −80°C following clarification and freeze-thaw treatment (41, 42). Cell debris was removed by centrifugation, and the virus-containing supernatant was passed through a 0.22-µm filter.

For amplification of infectious BKPyV stocks, WI-38 cells were infected at an MOI of 0.1. Culture supernatants were collected weekly, clarified by centrifugation, concentrated to one-tenth of the original volume, and stored at −80°C. At approximately 4 weeks post-infection, when pronounced cytopathic effects were observed, cells and supernatants were harvested and combined with supernatants collected at earlier time points. Virus-containing material was subjected to three freeze–thaw cycles, clarified by centrifugation, passed through a 0.22-µm filter, aliquoted, and stored at −80°C.

Infectious BKPyV titers were determined by focus-forming unit (FFU) assay. WI-38 cells were seeded at 3 × 10⁴ cells per well in 12-well plates and infected with serial dilutions of the virus stock (1:5, 1:10, 1:50, and 1:100). At 3 days post-infection (dpi), infected cells were detected by immunofluorescence staining for BKPyV VP1 and large T antigen (LT). Infected nuclei were quantified from three fields of view per dilution using 4 × 4 tile acquisition at 63× magnification, and infectious titers were expressed as FFU/mL.

SV40 stocks were propagated in CV-1 cells, and infectious titers were determined by fluorescent focus assay (40). MuPyV stocks were propagated in NMuMG cells, and infectious titers were determined by fluorescent focus assay using VP1 immunostaining. Titers were expressed as FFU/mL.

For BKPyV infection experiments, cells were seeded 16–24 h before infection and infected at approximately 50–80% confluence.

### Quantification of BKPyV genome copies

BKPyV genome copies were quantified by TaqMan qPCR targeting the VP1 region using the primers described in supplementary Table S1. For the secondary screening assay, qPCR was performed using the LightCycler 480 II system (Roche) and the QuantiFast Pathogen +IC Kit (Qiagen).

### Compounds

The primary screening library comprised approximately 28,000 small molecules assembled from four compound libraries obtained from Enamine and ChemDiv and provided by the German Center for Research in Infection, DZIF. The compounds had not previously been screened for antiviral activity against polyomaviruses.

Compound stocks were prepared in DMSO at 10 mM. Unless otherwise indicated, compounds were freshly diluted in the appropriate growth medium to a final concentration of 10 µM and added concomitantly with the virus inoculum. DMSO-treated infected cells served as vehicle controls.

Cidofovir (CDF) (MedChemExpress) and hexachlorophene (HXC) (Sigma-Aldrich) were evaluated during assay establishment as reference compounds.

### Phenotypic high-throughput screening using SV40-infected CV-1 reporter cells

A cell-based phenotypic high-throughput screening assay was performed using stably RFP-expressing CV-1 reporter cells and SV40 as a surrogate polyomavirus. The reporter cells were generated by lentiviral transduction of CV-1 cells with LeGO-EFS-RFP (43), followed by G418 selection and fluorescence-activated single-cell sorting. Three RFP-positive clones were subsequently pooled to establish the reporter-cell population used for high-throughput screening.

For the primary screen, CV-1 reporter cells were seeded at 4 × 10² cells per well in 384-well plates one day before infection, corresponding to approximately 25–40% confluence at the time of infection. Approximately 28,000 compounds were screened using a robotic BSL-2-compatible high-throughput platform at Hannover Medical School. Cells were infected with SV40 at an MOI of 0.5 and treated with individual compounds at a final concentration of 10 µM. Test compounds were analyzed in duplicate. DMSO-treated infected cells served as vehicle controls, uninfected reporter cells represented the maximal reporter-cell signal, and HXC-treated infected cells served as reference inhibitor controls.

A total of 243 384-well plates were analyzed. Each plate contained 52 uninfected control wells, 20 DMSO-treated infected control wells, 12 HXC-treated infected control wells, and 114 test compounds analyzed in duplicate.

At 6 days post-infection (dpi), culture medium was removed and cells were washed twice with DPBS. RFP fluorescence was quantified using a Cytation 5 Cell Imaging Multi-Mode Reader (BioTek). In parallel, automated wide-field fluorescence images were acquired using a 4× objective, with two fields captured per well. Mean RFP fluorescence intensity was determined from the acquired images using Gen5 Image+ 2.09 software (BioTek).

### Confirmatory screening of primary hits

The 98 compounds identified in the primary screen were reordered from Enamine or ChemDiv and subjected to confirmatory screening using the same experimental conditions as the primary HTS. Each compound was analyzed on six independent plates with two replicate wells per plate, resulting in 12 measurements per compound. Each plate additionally contained 20 HXC-treated infected reference wells, 36 DMSO-treated infected control wells, and 60 uninfected control wells.

### Secondary validation against BKPyV in primary RPTE cells

The 33 compounds confirmed in the SV40 assay were evaluated for antiviral activity against BKPyV in primary human RPTE cells. Cells at passage 4 were seeded in 96-well plates and either left uninfected or infected with BKPyV at an MOI of 0.5. Infected cells were treated with DMSO, HXC, CDF, or the respective test compound. At 6 dpi, genomic DNA was isolated from both cell-associated and culture-supernatant fractions and BKPyV genome copies were quantified by TaqMan qPCR targeting VP1.

### Concentration-response and cytotoxicity analyses

Concentration–response experiments were performed to determine the antiviral potency of compounds C1–C16 against BKPyV. Primary human RPTE cells were infected with BKPyV at an MOI of 0.5 and treated with increasing concentrations of the respective compounds, covering a concentration range of 0.01–100 µM. At 6 days post-infection (dpi), culture supernatants and cell-associated fractions were collected separately, genomic DNA was isolated, and BKPyV genome copies were quantified by qPCR. Viral genome copies were normalized to the corresponding DMSO-treated infected controls, which were set to 100%. Concentration–response experiments were performed in triplicate.

Concentration–response curves were fitted using GraphPad Prism version 11.0.2 (GraphPad Software), and IC₅₀ values were calculated independently for each compound and MOI.

For cytotoxicity analyses, uninfected primary RPTE cells were exposed to increasing concentrations of the respective compounds for 6 days. Cell viability was assessed using MTT (5mg/ml in DPBS4 h incubation, dissolution of formazan in isopropanol containing 0.04 M HCl, and absorbance at 570 nm with 630 nm as reference), and half-maximal cytotoxic concentrations (CC₅₀) were derived from the corresponding concentration–response curves. The selectivity index (SI) was calculated as: SI = CC₅₀ / IC₅₀

### In silico assessment of compounds

Compounds were evaluated according to their antiviral potency, cytotoxicity, selectivity, predicted toxicological liabilities, physicochemical properties, and chemical accessibility. Toxicological alerts were assessed using ProTox 3.0 DEREK (44). Molecular weight and cLogD at pH 7.4 were obtained using (45) and compliance with Lipinski criteria was evaluated (46).

### Antiviral activity against murine polyomavirus

To assess whether antiviral activity extended to murine polyomavirus (MuPyV), compounds C1–C16 were evaluated in NMuMG cells. Cells at passages 5–8 were seeded in 24-well plates and either left uninfected or infected with MuPyV at an MOI of 0.5. Infected cells were treated with DMSO, hexachlorophene (HXC), cidofovir (CDF), or the respective test compound at a final concentration of 10 µM. At 6 days post-infection (dpi), cell-associated material and cell-free culture supernatants were collected separately. Genomic DNA was isolated using the QIAamp DNA Mini Kit (Qiagen).

MuPyV genome copies were quantified by SYBR Green qPCR using the primers described in supplementary Table S1. qPCR was performed on a QuantStudio 3 Real-Time PCR System (Applied Biosystems) in a total reaction volume of 20 µL, containing 2 µL template DNA, 10 µL PerfeCTa SYBR Green FastMix (Quanta Bio), 0.8 µL of each primer, and 6.4 µL nuclease-free water. Cycling conditions consisted of an initial denaturation at 95°C for 10 min, followed by 40 cycles of 95°C for 15 s and 60°C for 30 s, followed by melting-curve analysis. Samples were analyzed in technical duplicate. Absolute MuPyV genome copy numbers were determined using a standard curve generated from serial dilutions of pBluescript-sk+PTA, containing the MuPyV genome, ranging from 10³ to 10⁸ genome copies/µL. Cell-associated MuPyV genome copies were normalized to mouse GAPDH where applicable, and viral loads were expressed relative to DMSO-treated infected controls

### Time-of-addition experiments

To determine the temporal window during which the compounds retained antiviral activity, time-of-addition experiments were performed in hTERT RPTE cells. Cells were infected with BKPyV at an MOI of 0.1, and compounds were added at a final concentration of 2-10µM at 0, 3, 6, 12, 24, 48, or 96 hpi.

Extracellular BKPyV DNA was quantified by qPCR at 6 dpi and normalized to the corresponding DMSO-treated infected controls.

### Compound pretreatment

To assess whether compound exposure prior to infection affected subsequent BKPyV replication, hTERT RPTE cells were pretreated with compounds C1–C16 at 2-10µM for 3hrs. Following pretreatment, compound-containing medium was removed, cells were washed 1x in DPBS and fresh compound-free medium was added. Cells were subsequently infected with BKPyV at an MOI of 0.5 and maintained in the absence of compound. At 6 dpi, culture supernatants were collected and extracellular BKPyV genome copies were quantified by qPCR as described above. Viral loads were normalized to DMSO-pretreated infected controls.

### MOI-dependent concentration–response analysis

To assess the effect of viral inoculum on antiviral potency, concentration–response analyses were performed for C5, C8, and C9 at BKPyV MOIs of 0.05, 0.5, and 5. hTERT RPTE cells were seeded one day before infection and infected with BKPyV at the indicated MOI. Cells were treated with increasing concentrations of the respective compounds (0.1, 0.25, 0.5, 0.75, 1.25, 2.5, 5, 10, and 30 µM). DMSO-treated infected cells at the corresponding MOI served as vehicle controls.

At 6 days post-infection (dpi), culture supernatants were collected and extracellular BKPyV genome copies were quantified by qPCR as described above. Viral genome copies were normalized to the corresponding DMSO-treated infected controls. Concentration–response curves were fitted using GraphPad Prism version 11.0.2 (GraphPad Software), and IC₅₀ values were calculated independently for each compound and MOI.

### BKPyV VLP production

BKPyV virus-like particles (VLPs) were produced essentially as described previously using the protocol developed by the Buck laboratory (47). HEK293 cells were transfected with the BKPyV capsid expression plasmids pWB2b (VP1/VP2) (Addgene plasmid #32094) and pWB3b (VP1/VP3) (Addgene plasmid #32106). Following capsid expression, cells were harvested and lysed, and VLP-containing lysates were subjected to nuclease treatment and capsid maturation. VLPs were purified by iodixanol (OptiPrep) density-gradient ultracentrifugation, and VLP-containing fractions were collected. Purified VLPs were fluorescently labelled using an Alexa Fluor 488 Protein Labeling Kit (Thermo Fisher Scientific, A10235) according to the manufacturer’s instructions and subsequently used for VLP attachment experiments.

### BKPyV virus-like particle attachment assay

To determine whether compounds affected BKPyV attachment to the cell surface, 7 × 10⁵ hTERT RPTE cells per well were seeded in 6-well plates and treated with the respective compounds or an equivalent concentration of DMSO. At 24 h after treatment, cells were detached using CellStripper non-enzymatic cell dissociation buffer (Corning) to preserve cell-surface receptor integrity. Cells were incubated with CellStripper for a maximum of 20 min, collected by centrifugation at 452 × g for 5.5 min at 4°C, and washed once with DPBS.

Cells were subsequently incubated with Alexa Fluor 488-labelled BKPyV virus-like particles (VLPs) at an MOI of 1 in a total volume of 500 µL for 2 h at 4°C under gentle agitation. Incubation at 4°C was used to permit particle attachment while minimizing internalization. Following VLP exposure, cells were washed with DPBS to remove unbound particles and fixed with 4% paraformaldehyde (PFA) for 10 min on ice. After fixation, cells were washed once with DPBS and resuspended in DPBS containing 1% FBS for flow-cytometric analysis.

For each sample, 10,000 events were acquired (LSR Fortessa). Doublets and cellular debris were excluded based on forward- and side-scatter characteristics, and cell-associated Alexa Fluor 488 fluorescence was quantified as a measure of VLP attachment (FACSDiva 8.0.1).

### Analysis of BKPyV transcription

To determine the effects of C5, C8, and C9 on viral transcription, hTERT RPTE cells were infected with BKPyV and treated with the respective compound or an equivalent concentration of DMSO. At 48 hrs pi, total RNA was isolated (AllPrep DNA/RNA/Protein Mini Kit, Qiagen, Germany)and treated with DNA-free DNase (Gibco) to remove residual genomic DNA.

For cDNA synthesis, 1–5 µg total RNA was incubated with random hexamer primers for 5 min at 65°C and immediately cooled on ice for 2 min. Reverse transcription was performed using SuperScript IV reverse transcriptase in the presence of 5× SuperScript IV buffer, dNTPs, DTT, murine RNase inhibitor, and MgCl₂. Reactions were incubated for 10 min at 23°C followed by 30 min at 50°C. Minus-reverse-transcriptase controls were included to monitor residual genomic DNA contamination.

BKPyV transcript abundance was quantified by SYBR Green RT-qPCR targeting the early transcripts large T antigen (LT) and small T antigen (sT) and the late transcript VP1. Primers were designed to preferentially span exon–exon junctions or exon–intron boundaries to minimize amplification from residual viral genomic DNA. Expression of the cellular housekeeping genes GAPDH and TBP was determined in parallel by multiplex TaqMan assays and used for normalization. Each experimental condition was analyzed in technical triplicate, and amplification specificity of the SYBR Green assays was confirmed by melt-curve analysis.

Relative viral transcript abundance was calculated using the comparative Ct (2⁻ΔΔCt) method, with viral transcript levels first normalized to the housekeeping genes and subsequently expressed relative to DMSO-treated BKPyV-infected cells. Primer sequences are provided in supplementary Table S1.

### RNA isolation and transcriptomic profiling

For transcriptomic profiling, hTERT RPTE cells were analyzed 48 hrs p.i. and/or compound treatment. Experimental conditions comprised non-infected and BKPyV-infected cells treated with DMSO, C5, C8, or C9. Two independent biological replicates were analyzed for each condition.

High RNA integrity was confirmed by applying a TapeStation RNA ScreenTape analysis (Agilent, 5067-5576, 5067-5577, 5067-5578). Per sample, 1 µg total RNA (quantified by Qubit RNA HS Assay, Thermo Fisher, Q32852) was Poly(A)-captured applying the Lexogen Poly(A) RNA Selection Kit and further processed via the RNA-Seq V2 Library Prep Kit with UDIs (Lexogen, catalog number: 181.96) in the long insert size variant (RTL) according to manufacturer’s instructions (applying 11 cycles of Library Amplification PCR, step 6.3, User Guide version 171UG394V0101).

Libraries were quality controlled on a TapeStation D5000 Assay (Agilent, D5000 ScreenTape 5067-5588 with D5000 Reagents 5067-5589) and were sequenced on an Element Biosciences AVITI instrument (2×150 Sequencing Kit Cloudbreak Freestyle, product number: 860-00012) in a paired-end mode. After demultiplexing via bases2fastq, all samples passed quality control by FastQC and MultiQC (48) and were subjected.

### Transcriptome analysis

Gene abundance for each sample were quantified with human gene annotations from gencode (version 49) for GRCh38 genome assembly using salmon (v1.10.0) (49), and imported using R package tximport (50). Counts normalization and multi-factor differential expression (DE) analysis was performed using DESeq2 package (51). Null variance of Wald test statistic output by DESeq2 was re-estimated using R package fdrtool (52) to calculate p-values (and adjusted using Benjamini-Hochburg method) for the final list of differentially expressed gene. FDR (BH-adjusted p-values) < XX was used a criteria for the final DE gene list. Volcano plot were created using EnhancedVolcano R package [5] and Gene ontology enrichment analysis was performed using R package clusterProfiler (53).

### Differential gene expression and functional enrichment analysis

Differential gene expression analysis was performed using DESeq2. For definition of the BKPyV infection-associated transcriptional signature, BKPyV-infected, DMSO-treated hTERT RPTE cells were compared with non-infected, DMSO-treated cells at 48 h post-infection. Compound-associated transcriptional responses were determined by comparing C5-, C8-, or C9-treated cells with the corresponding DMSO-treated controls separately in non-infected and BKPyV-infected cells. Two biological replicates were included per experimental condition.

Principal component analysis (PCA) was performed on normalized gene-expression data to assess global transcriptional differences between experimental conditions. Genes with an adjusted p value < 0.05 and log₂ fold change ≥ 1 were considered differentially expressed. Gene Ontology (GO) enrichment analysis was performed on DEGs to identify overrepresented functional categories, including Biological Process and Cellular Component terms. Enrichment significance was assessed using Fisher’s exact test. For visualization, enriched GO terms were ranked according to statistical significance, and selected genes contributing to functionally relevant categories were displayed according to their log₂ fold changes and adjusted p values.

### Gene set enrichment analysis

Genes were ranked according to the DESeq2 Wald statistic and subjected to gene set enrichment analysis (GSEA) using the fgsea multilevel algorithm. Gene sets were obtained from the Hallmark collection of the Molecular Signatures Database (MSigDB). Pathways were considered for further analysis using an exploratory threshold of adjusted p < 0.1, an absolute normalized enrichment score (|NES| > 1.5), and a minimum leading-edge size of >10 genes. For visualization, the ten enriched pathways with the highest absolute NES values were selected. For each pathway, the five leading-edge genes with the largest absolute log₂ fold changes were extracted and displayed.

### Statistical analysis

Statistical analyses and graphical presentation of experimental data were performed using GraphPad Prism version 11.0.2 (GraphPad Software, Boston, MA, USA) unless otherwise indicated. Data are presented as mean ± standard deviation (SD) or as individual values, as specified in the respective figure legends. The number of independent biological replicates (n) and, where applicable, technical replicates are indicated in the corresponding figure legends.

For experiments in which viral genome copies or transcript levels were expressed relative to a control condition, values were normalized to the corresponding DMSO-treated control as described for the respective assay. Statistical comparisons were performed using two-way Anova followed by Dunnett’s multiple comparisons test versus DMSO. *p < 0.05; **p < 0.01; ***p < 0.001, or one sample T-test against a hypothetical value of 100% or two-tailed Student’s t test Error bars represent means ± SEM of biological replicates. *, P < 0.05; **, P < 0.01; ***, P < 0.001, with correction for multiple comparisons where applicable. A p value <was considered statistically significant.

## Data Availability statement

All data supporting the findings of this study are available within the article and its supplemental material. Additional materials and data are available from the corresponding author upon reasonable request.

## Acknowledgement

We thank Kerstin Reumann for technical assistance with NGS library preparation and Chris Sullivan (University of Texas at Austin, Austin, TX, USA) for providing resources and support for the MuPyV experiments.

This work was supported by the Deutsche Forschungsgemeinschaft (DFG, German Research Foundation) -GRK2771 – project no. 453548970 and Project-ID 443644894 – Research Unit FOR 5200 (DEEP-DV). This work was in parts supported by the German Centre for Infection Research (DZIF, grant number TTU 07.828)

## Author contribution

**Clara Husser:** Conceptualization, Methodology, Investigation, Formal analysis, Data curation, Visualization, Writing – original draft, Writing – review & editing.

**Hannes Roggenkamp:** Investigation, Methodology, Visualization.

**Emma Kraus:** Investigation, Methodology.

**Claudia Schmidt:** Investigation, Methodology.

**Jessica Rückert:** Investigation, Methodology.

**Patrick Blümke:** Investigation, Formal analysis.

**Sanamjeet Virdi:** Formal analysis, Data curation, Visualization.

**Thomas Schulz:** Conceptualization, Funding acquisition, Resources.

**Adam Grundhoff:** Conceptualization, Methodology, Funding acquisition, Resources, Writing.

**Nicole Fischer:** Conceptualization, Methodology, Supervision, Project administration, Funding acquisition, Resources, Writing – review & editing.

## Supplementary Materials

Supplementary Figures S1-S5

Supplementary Table S1

## Supplementary Figure Legends

**Supplementary Figure S1. Validation of hexachlorophene as a positive control for the SV40 phenotypic screening assay.**

**(A)** Viability of CV-1 cells treated with 10 µM hexachlorophene (HXC) or cidofovir (CDF), added at the time of infection, was assessed by MTT assay after 6 days of culture. Absorbance values were normalized to uninfected, DMSO-treated cells, which were set to 100%. (B, C) SV40 infection was assessed at 6 days post-infection (p.i.) by quantification of RFP fluorescence intensity (FI) in reporter cells using a multimode microplate reader **(B)** or automated wide-field fluorescence microscopy (BioTek) **(C).** Infected reporter cells were treated with DMSO, HXC, or CDF at the time of infection. RFP fluorescence values were normalized to uninfected reporter cells, which were set to 100%. HXC produced a robust inhibitory phenotype across both fluorescence-based readouts and was therefore selected as the positive control for the subsequent screening.

**Supplementary Figure S2**: **Representative plates showing compounds that inhibit SV40 progeny production and cytopathic effect (CPE) formation in CV-1 reporter cells**. Hit selection was performed on a plate-by-plate basis using two independent cut-offs: relative RFP fluorescence intensity (FI) >20% or fold change relative FI >2. Representative data from three independent plates are shown. Data sets used for hit selection based on the relative FI cut-off are shown in (A–C), whereas data sets used for hit selection based on fold change relative FI are shown in (D–F). Relative FI values required for the first cut-off and fold change relative FI values required for the second cut-off for the top 98 selected compounds, as measured by plate reader, are shown in (G), with corresponding imaging-based data shown in (H).

**Supplementary Figure S3. Activity of selected BKPyV inhibitors against murine polyomavirus (MuPyV).**

**(A–B)** NMuMG cells were infected with MuPyV at a multiplicity of infection (MOI) of 0.5 and treated with DMSO, hexachlorophene (HXC), cidofovir (CDF), or compounds C1–C16 at 10 µM. At 6 days post-infection (p.i.), MuPyV genome copies were quantified by qPCR in culture supernatants **(A)** and cell-associated fractions **(B)**. Viral genome copies were normalized to those of DMSO-treated infected cells, which were set to 100%.

**Supplementary Figure S4. Differential gene expression associated with BKPyV infection in hTERT RPTE cells.**

Transcriptomic profiling was performed in hTERT RPTE cells 48 h post-infection by comparing BKPyV-infected with non-infected control cells, both maintained in the presence of DMSO. **(A)** Principal component analysis (PCA) of the transcriptomic profiles of BKPyV-infected and non-infected cells. Each point represents an individual biological replicate. The first and second principal components (PC1 and PC2) account for 61.5% and 20.3% of the total variance, respectively, and show separation of BKPyV-infected from non-infected samples. **(B)** Volcano plot showing differential gene expression between BKPyV-infected and non-infected cells. The x-axis represents log2 fold change and the y-axis represents −log10(adjusted p value). Significantly upregulated and downregulated genes at an FDR of 0.05 are shown in red and blue, respectively, whereas genes not meeting the significance criteria are shown in black. A total of 473 genes were upregulated and 301 genes were downregulated in BKPyV-infected relative to non-infected cells. For visualization, the y-axis was capped at 40; adjusted p values exceeding this scale are not displayed at their full magnitude.

**Supplementary Figure S5: Transcriptomic characterization of C5-treated non-infected and BKPyV-infected hTERT RPTE cells.**

Transcriptomic profiling was performed in hTERT RPTE cells 48 h post-infection to characterize C5-induced transcriptional changes in non-infected (NI) and BKPyV-infected cells relative to the corresponding DMSO-treated controls. **(A)** Principal component analysis (PCA) of DMSO- and C5-treated non-infected and BKPyV-infected cells. Each point represents an individual biological replicate. PC1 and PC2 account for 36.1% and 20.1% of the total variance, respectively. **(B)** Venn diagram showing the overlap between the BKPyV infection-associated transcriptional signature and the transcriptional responses to C5 treatment in non-infected (C5 (NI)) and BKPyV-infected (C5 (BKPyV)) cells. Numbers indicate genes unique to or shared between the respective signatures. **(C)** Number of differentially expressed genes identified for the indicated transcriptional signatures. **(D)** Functional enrichment analysis of the C5-induced transcriptional response, showing enriched pathways related predominantly to translation, mitochondrial translation, tRNA aminoacylation, and transcriptional regulation. Bars indicate the number of genes associated with each term, and color represents −log10(adjusted p value). **(E–F)** Gene Ontology Cellular Component enrichment analysis of the C5-induced transcriptional response in non-infected **(E)** and BKPyV-infected **(F)** cells. Enriched cellular components are plotted according to fold enrichment. Dot size represents the number of genes associated with each term, and dot color indicates −log10(adjusted p value).

**Supplementary Table S1:**
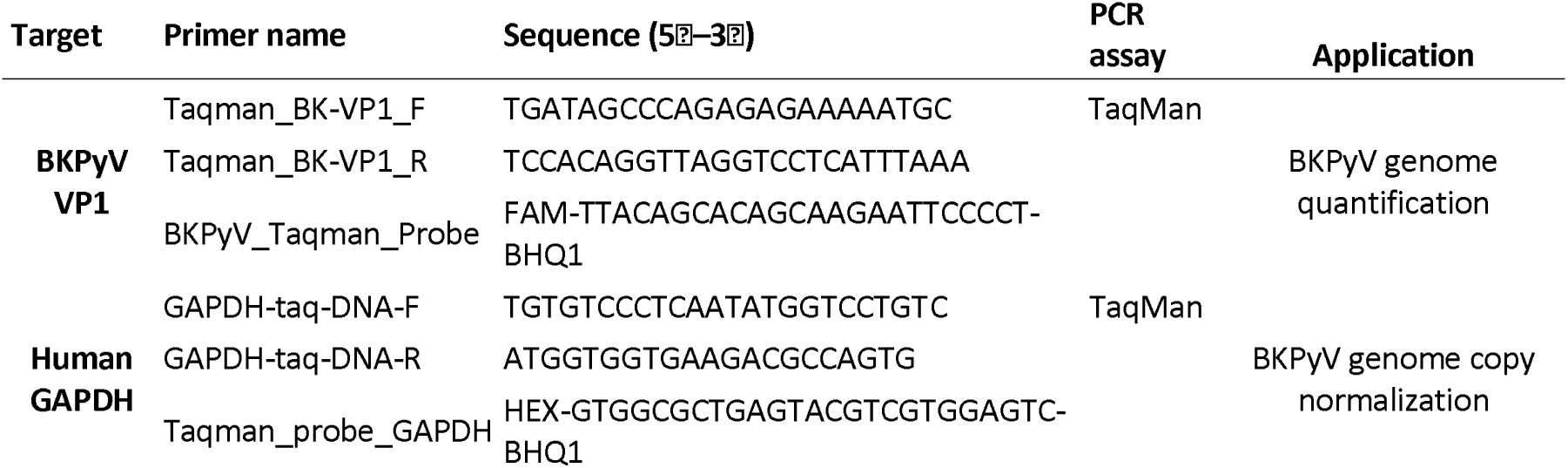
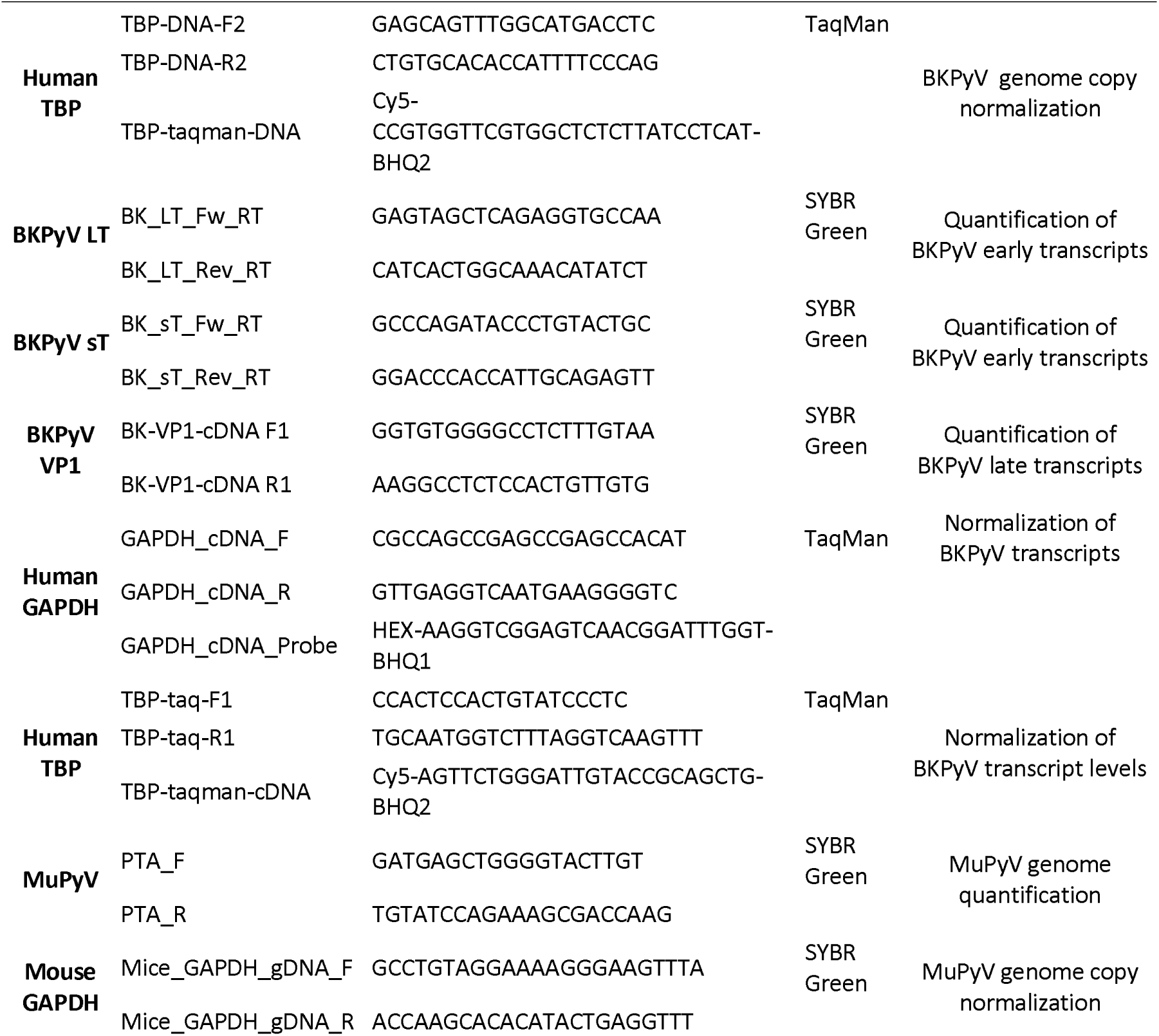
Primers and probes for qPCR and RT-PCR analyses.

## Notes

### Competing Interest Statement

The authors have declared no competing interest.

