## Supplementary Figures for "Phenotypic Screening Identifies Small-Molecule Inhibitors with Distinct Activities across the BK Polyomavirus Life Cycle"

### Supplementary Figure S1

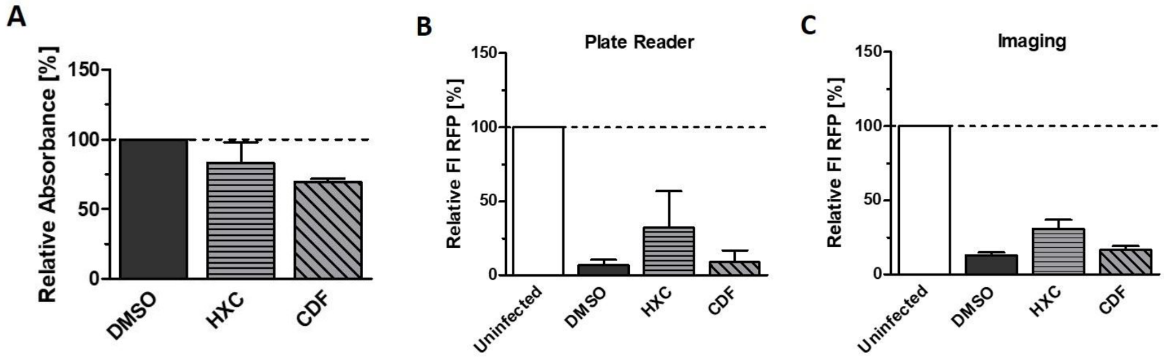

#### Supplementary Fig. S2

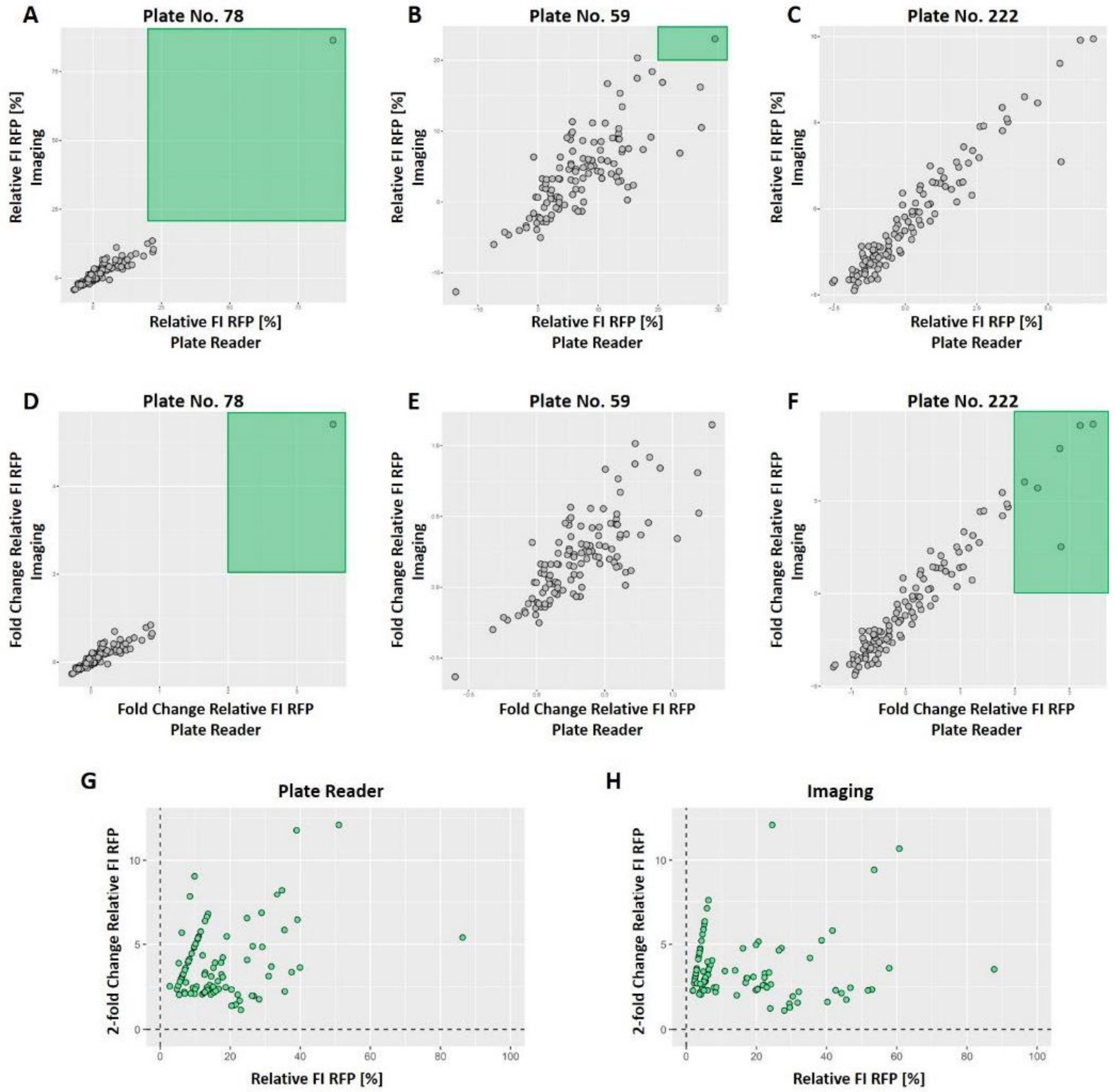

Suppl. Fig S3

A

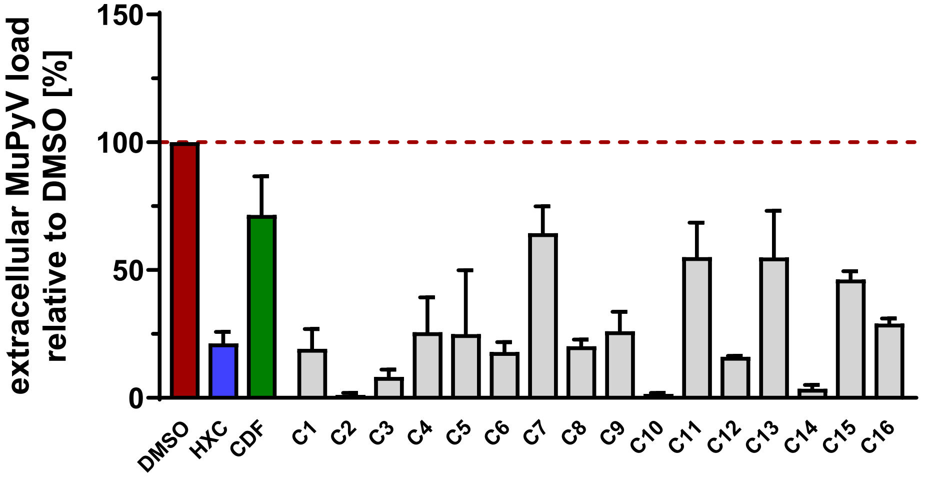


B

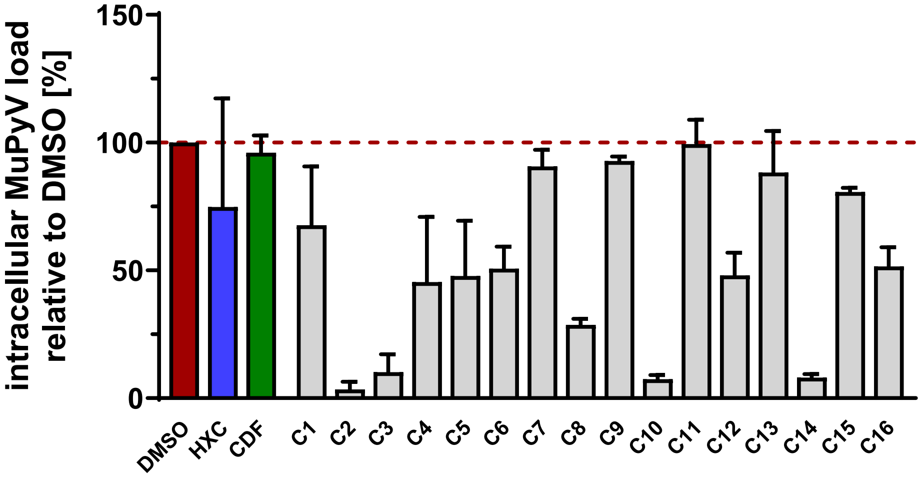

**Suppl. Fig S4**

**A**

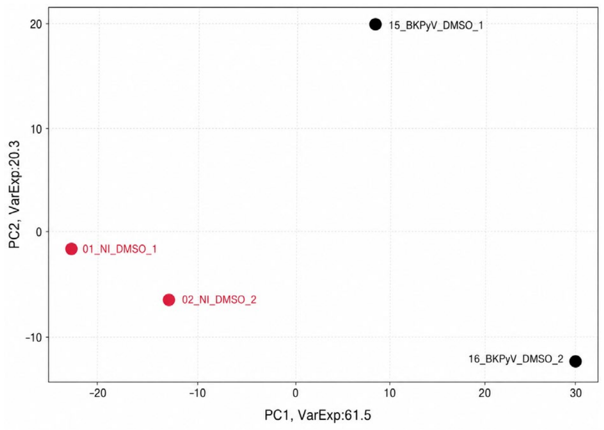


**B**

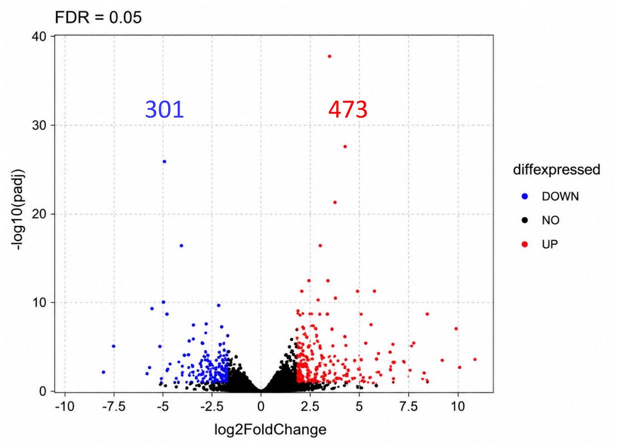


Y-axis capped at 40. Points above 40 not shown (see data: max -log10(padj) = 159.8)

**Suppl. Fig.S5**

A

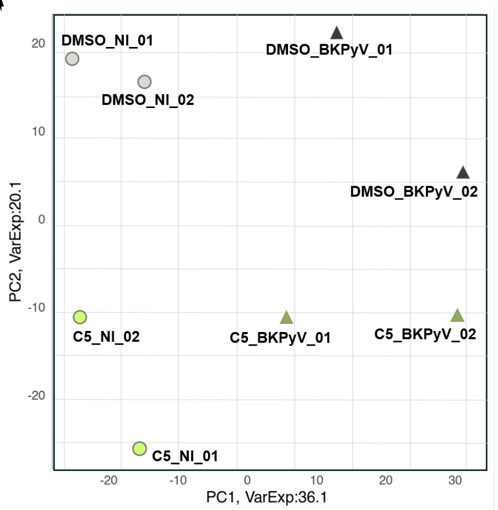


B

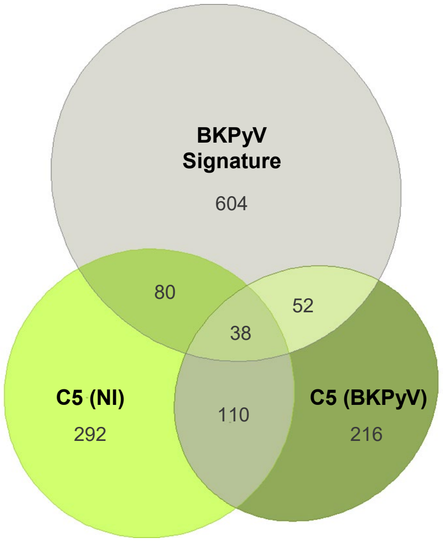


C

| | C5 (NI) | C5 (BKPyV) | BKPyV Signature |
|---|---|---|---|
| **UP** | 285 | 247 | 473 |
| **DOWN** | 235 | 169 | 301 |
| **Total** | 520 | 416 | 774 |

D

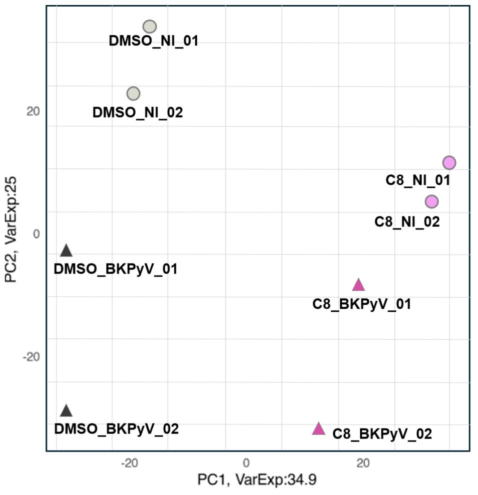


E

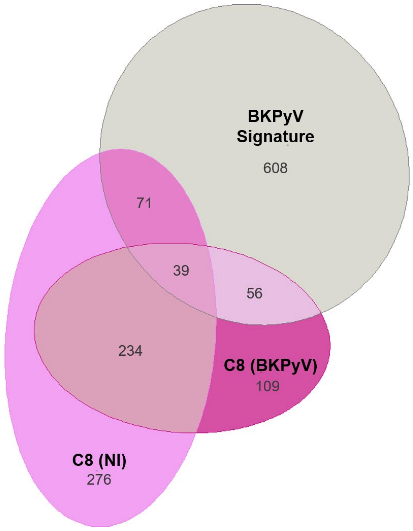


F

| | C8 (NI) | C8 (BKPyV) | BKPyV Signature |
|---|---|---|---|
| **UP** | 319 | 230 | 473 |
| **DOWN** | 301 | 208 | 301 |
| **Total** | 620 | 438 | 774 |
